# *In vivo* functional genomics identifies essentiality of potassium homeostasis in medulloblastoma

**DOI:** 10.1101/2022.07.23.501234

**Authors:** Jerry J. Fan, Xin Wang, Anders W. Erickson, Patryk Skowron, Xian Wang, Xin Chen, Guanqiao Shan, Shahrzad Bahrampour, Yi Xiong, Weifan Dong, Namal Abeysundara, Michelle A. Francisco, Ronwell J. Pusong, Raúl A. Suárez, Hamza Farooq, Borja L. Holgado, Xiaochong Wu, Craig Daniels, Adam J. Dupuy, Juan Cadiñanos, Allan Bradley, Anindya Bagchi, Branden S. Moriarity, David A. Largaespada, A. Sorana Morrissy, Vijay Ramaswamy, Stephen C. Mack, Livia Garzia, Peter B. Dirks, Siyi Wanggou, Xuejun Li, Yu Sun, Michael D. Taylor, Xi Huang

## Abstract

The identification of cancer maintenance genes—driver genes essential to tumor survival—is fundamental for developing effective cancer therapy. Transposon-based insertional mutagenesis screens can identify cancer driver genes broadly but not discriminate maintenance from progression or initiation drivers, which contribute to cancer phenotypes and tumorigenesis, respectively. We engineered a nested, double-jumping transposon system to first dysregulate gene expression during tumorigenesis and then restore gene expression following tumor induction, allowing for genome-wide screening of maintenance essentiality *in vivo*. In a mouse model of medulloblastoma, the most common pediatric malignancy, insertion and remobilization of this nested transposon uncovers potassium channel genes as recurrent maintenance drivers. In human medulloblastoma, KCNB2 is the most overexpressed potassium channel across Group 3, Group 4, and SHH subgroups, and *Kcnb2* knockout in mice diminishes the replicative potential of medulloblastoma-propagating cells to mitigate tumor growth. Kcnb2 governs potassium homeostasis to regulate plasma membrane tension-gated EGFR signaling, which drives proliferative expansion of medulloblastoma-propagating cells. Thus, our novel transposon system reveals potassium homeostasis as essential to tumor maintenance through biomechanical modulation of membrane signaling.

## INTRODUCTION

Continued mutagenesis during clonal expansion in cancer yields new subclones with distinct mutational landscapes that contribute to tumor progression and maintenance. The ability to distinguish cancer maintenance drivers, which are required to sustain tumor growth, from ineffective targets (initiation, progression, and passenger events) is critical for the development of cancer therapeutics. While insertional mutagenesis systems, such as Sleeping Beauty (SB) and piggyBac (PB), can discern driver from passenger events by mobilizing transposons randomly across the genome to cause gain- or loss-of-function mutations at target sites, they cannot further categorize cancer drivers^1–11^. Among these, initiation drivers confer tumor-forming capacity to normal cells, and progression drivers enhance cancer phenotypes, but neither are essential for clonal survival. Only maintenance drivers are necessary for tumor survival.

Identification of maintenance driver genes is highly relevant in medulloblastoma (MB), the most common, malignant, solid, pediatric cancer^12^. Current treatments, while effective in some MB subgroups, are non-targeted and bear debilitating long-term sequelae^13, 14^. Given the availability of tractable genetic mouse models for Sonic Hedgehog (SHH) MB, which comprises 30% of human MB cases, and the lack of effective targeted therapy, we engineered a novel transposon system to discover *bona fide* tumor maintenance genes in SHH MB^15–19^.

## RESULTS

### Insertional mutagenesis implicates potassium channels in medulloblastoma maintenance

To identify cancer maintenance drivers, we engineered an *in vivo* screening system using Lazy Piggy (LP), a hybrid transposon containing both SB and PB excision sequences flanking cargo capable of dysregulating gene expression. The LP transposon was created by nesting PB inverted terminal repeats (ITRs) medial to the SB ITRs of the T2/Onc2 transposon (**Figures 1A-C**). Depending on the transposon integration site, target gene expression can be activated or disrupted through murine stem cell virus (MSCV) promoter-splice donor (SD) elements and splice acceptor-polyadenylation signal (SA-pA) elements, respectively^2, 20^. SB transposase enables primary mobilization of the LP transposon from a donor concatemer into random genomic reintegration sites, with relocated transposons dysregulating gene expression. Subsequent activation of tamoxifen-inducible PB remobilizes and depletes insertion events by extracting the transposon and restoring normal gene function (**Figures S1A-B**). We spatially and temporally controlled the LP system by tissue-specific SB transposase expression and tamoxifen-inducible PB transposase, respectively. SB transposase mobilizes the hybrid LP transposon from chromosomal concatemers, with relocated transposons disrupting gene expression in cell lineages of interest. Subsequent activation of tamoxifen-inducible PB remobilizes and depletes insertion events by extracting the transposon, thereby restoring original gene function (**Figure 1B**). Initiation, passenger, and progression insertions in tumor cells are removed with no deleterious effect to clonal survival. In contrast, transposon remobilization from maintenance insertions mitigates tumor maintenance. This process depletes insertion events that are non-essential for clonal survival, while enriching for maintenance driver insertions to enable the identification of optimal targets for therapy development (**Figure 1C**).

**Figure 1.**
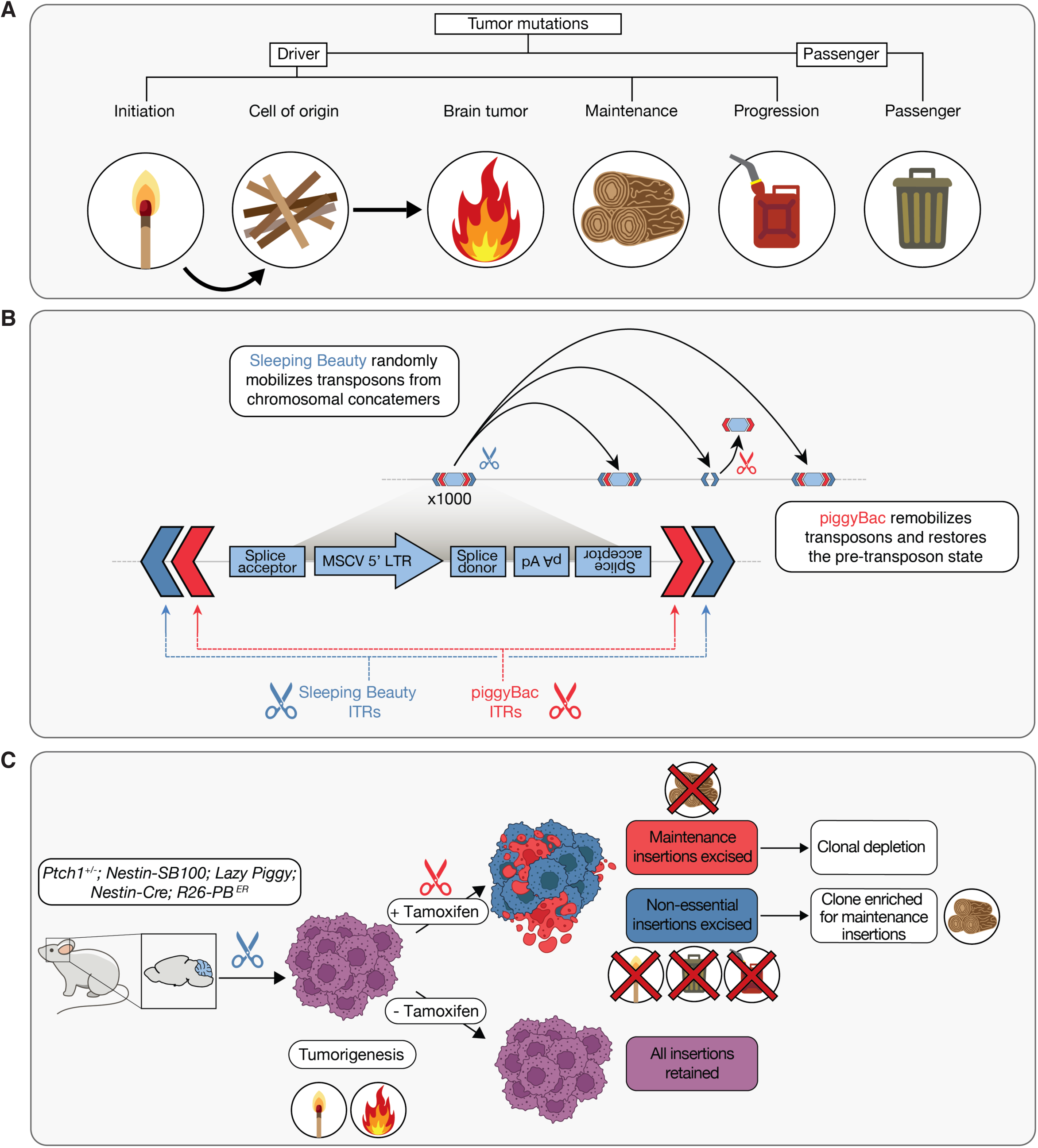
The Lazy Piggy transposon system enriches for cancer maintenance drivers. (A) Cartoon illustrating the rationale and design of the Lazy Piggy (LP) transposon system. Tumors harbor myriad mutations and genetic alterations, which can be classified as drivers and passengers (garbage). Drivers encompass initiation (match), maintenance (wood), and progression (gasoline) events. Initiating drivers transform non-tumor cells of origin to initiate tumorigenesis, maintenance drivers sustain continued tumor growth, and progression drivers elevate the malignant or metastatic potential of the tumor. Only targeting maintenance drivers will cause clonal collapse and tumor regression. (B) First, Sleeping Beauty (SB)-induced insertional mutagenesis drives MB progression. Second, low-dose tamoxifen induces PB-mediated remobilization, restoring original gene function of select insertion events. (C) While insertions at initiator, passenger, and progression genes are depleted with no consequence on tumor survival, transposon remobilization from maintenance drivers results in clonal collapse and depletion. Thus, this strategy enriches for retention of insertions at maintenance drives.

To identify genes involved in MB maintenance, we generated a LP-driven murine model of SHH MB (**Figures S1C-H**). Mice heterozygous for *Ptch1*, a negative regulator of the Shh pathway, sporadically develop MB with low latency^15^. In *Ptch1*^+/–^ mice, LP mutagenesis is lineage-restricted to neural progenitors of the developing mouse cerebellum by Nestin-driven SB transposase (*Nestin:Luc-SB100*) and temporally regulated by tamoxifen-inducible PB transposase (*Nestin-Cre; R26-LSL-mPB-ER^T^*^2^). To account for donor chromosome insertion bias, we generated separate founder mice harboring LP concatemers on chromosomes 7 and 10 (**Figures S1C-D**).

LP mutagenesis generated a highly penetrant model of SHH MB, accelerating medulloblastomagenesis in quintuple transgenic mice (*Ptch1^+/-^; Nestin:Luc-SB100^+/-^; Lazy Piggy^+/-^; Nestin-Cre^+/-^; R26-LSL-mPB^ERT2+/-^*) compared to *Ptch1*^+/–^ mice or quadruple transgenic mice without the *Ptch1*^+/–^ allele (*Nestin:Luc-SB100^+/-^; Lazy Piggy^+/-^; Nestin-Cre^+/-^; R26-LSL-mPB^ERT2+/-^*) (**Figure 2A**). Following tumor formation, low-dose tamoxifen partially depletes primary insertions by remobilizing cargo and restoring normal gene function. Remobilization of tumor-essential (i.e. maintenance) insertions leads to clonal depletion. Remobilization of initiator, progressor, and passenger insertions does not impact clonal survival and leads to enrichment of maintenance insertions. Transposon remobilization by low-dose tamoxifen does not alter the survival of quintuple mice (**Figure S2G**), as the small proportion of remobilizations depletes non-essential insertions and is not curative. We performed restriction-splink PCR^21, 22^ and gene-centric common insertion site (gCIS) analysis^23^ to determine reintegration sites with and without secondary mobilization in tumors from quintuple transgenic mice with (TAM^+^) or without tamoxifen (TAM^-^) treatment (**Figures S2A-C**). Insertions in two potassium channel genes, *Kcnb1* and *Kcnh2,* were enriched in TAM^+^ mice (**Figures 2B** and **S2D-E**). Additionally, we performed bulk transcriptomic analysis of TAM^+^ and TAM^-^ tumors to identify overexpressed genes in tumors enriched for maintenance insertions. Bulk RNA sequencing (RNA-seq) also revealed upregulated expression of additional potassium channel genes in TAM^+^ tumors (**Figures 2C** and **S2F**). Thus, the LP screen implicates potassium channels as potential regulators of MB maintenance.

**Figure 2.**
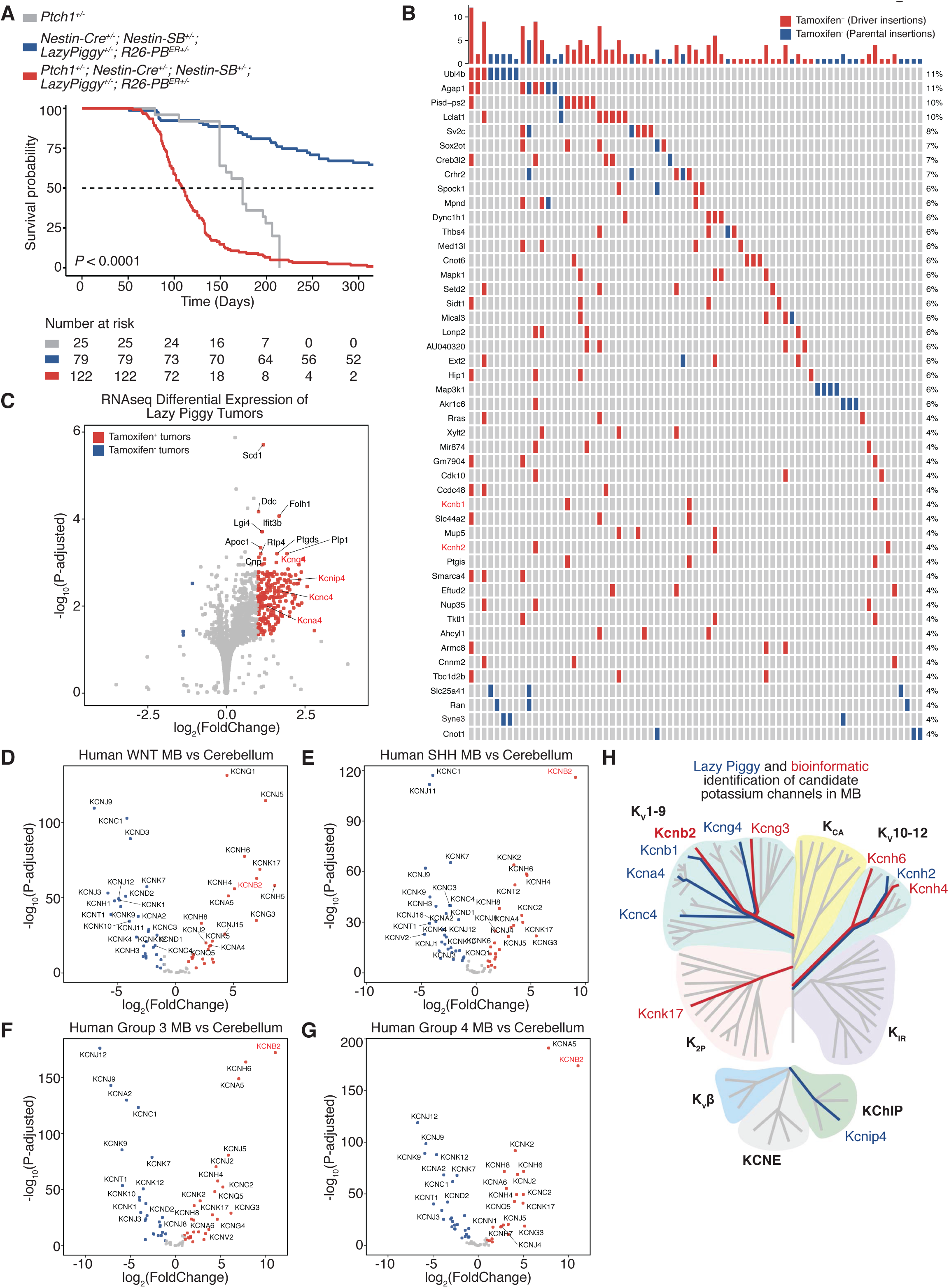
Lazy Piggy screening implicates potassium channels in medulloblastoma maintenance. (A) *Ptch1^+/–^* mice with LP transposition experienced poorer survival compared to *Ptch1^+/–^* mice or mice with LP transposition alone. (B) Overview of the significant gCIS insertions in candidate genes from all sequenced samples (n = 62) comprised of tamoxifen-negative tumors with ‘parental insertions’ that have not been remobilized, and tamoxifen-treated tumors with maintenance ‘driver insertions’ enriched by piggyBac-mediated transposon remobilization. (C) RNA-seq differential expression of tamoxifen-treated versus untreated tumors. Pink dots represent genes upregulated in tamoxifen-treated tumors that have been enriched for maintenance insertions. Outlier gene *Scd2* omitted for visualization (log2FC 0.49; padj 1e-11). (D-G) KCNB2 is upregulated among potassium channels in published data across human WNT (n = 56), SHH (n = 182), Group 3 (n = 131), and Group 4 medulloblastoma (n = 289) compared to human control cerebellum (n = 9) ^58^. Significantly differentially expressed genes are indicated in color (log2fold change > 1, Padj < 0.05). (H) Potassium channel phylogeny (gray lines) illustrating potassium channel genes identified from the Lazy Piggy screen (blue lines) and the most over-expressed potassium channel from patient expression analysis (red line). Bolded text indicates different classes of potassium channel genes (Kv, K2P, KCA, KIR, KVβ, KCNE, KChIP).

While the membrane localization and availability of pharmacological agents makes potassium channels attractive therapeutic targets^24–26^, it is critical to identify potassium channel targets with MB-specific function. *Kcnb1* knockout mice are hyperactive and prone to seizures^27^. *KCNB1* mutations are implicated in epileptic encephalopathy^28^ and neurodevelopmental disorders^29^ in humans. KCNH2 is a critical mediator of cardiac action potentials and its dysfunction can result in fatal cardiac arrhythmias^30^. Given the important functions of KCNB1 and KCNH2 in normal physiology, we examined potassium channel expression in MB more broadly to identify additional targets. Differential gene expression analysis of RNA-seq data from human MB and cerebellum^31^ identified *KCNB2,* the human paralog of *Kcnb1*, as the top-ranked potassium channel which is at least five-fold overexpressed across all four MB subgroups. Furthermore, *KCNB2* is the most highly expressed potassium channel across SHH, Group 3, and Group 4 MB, the three subgroups with poorest prognosis which collectively account for ∼90% of all MB (**Figure 2D-G**). While KCNB1 and KCNB2 share structural and functional similarities (**Figure 2H**), KCNB2 is not implicated in human disorders^29^. Taken together, our integrated cross-species study leverages both LP functional genomics to highlight potassium channels as MB maintenance drivers and orthogonal expression analysis to identify KCNB2 as the potassium channel with highest potential for MB-specificity. Therefore, we investigated the function of KCNB2 in MB.

### Kcnb2 is dispensable during normal mouse development

We first studied Kcnb2 knockout mice to determine its role during normal mouse development (**Figure 3A-F** and **S3A-F**). Kcnb2 knockout mice are viable (**Figure 3A**), fertile (**Figure 3D**), produced offspring at expected mendelian ratios (**Figure 3B**) and displayed no defects in development (**Figure S3A**), body weight at weaning (**Figure 3C**), or brain morphology (**Figure 3E** and **S3B**). We observed no overt differences in the cerebellar architecture of Kcnb2 knockout mice at postnatal day 7 and 30 (P7 and P30) (**Figure 3F** and **S3C-F**). These data indicate that Kcnb2 is non-essential for normal mouse development.

**Figure 3.**
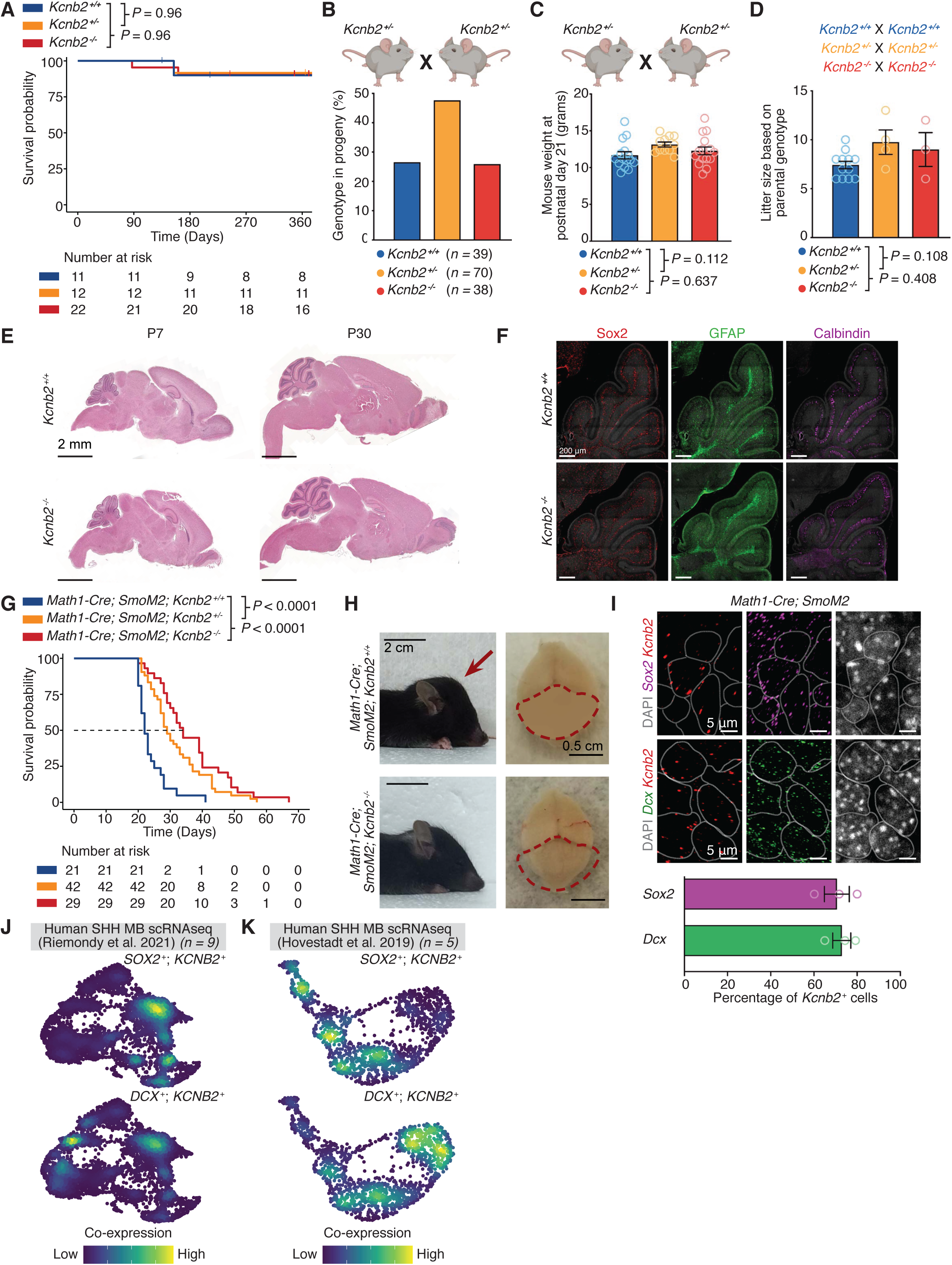
Kcnb2 is a medulloblastoma-specific vulnerability. (A) Kaplan-Meier survival plot of *Kcnb2^+/+^*, *Kcnb2^+/-^*, and *Kcnb2^-/-^* mice. (B) Genotypes of offspring obtained from crossing *Kcnb2^+/-^* parents. (C) Weight of *Kcnb2^+/+^*, *Kcnb2^+/-^*, and *Kcnb2^-/-^* mice at postnatal day 21. (D) Litter sizes obtained from breeding pairs of *Kcnb2^+/+^*, *Kcnb2^+/-^*, and *Kcnb2^-/-^* mice. (E) Representative histology of sagittal brain sections from P7 and P30 mice. (F) Representative immunohistochemistry of Sox2, GFAP, and Calbindin from sagittal cerebellum sections of *Kcnb2^+/+^* and *Kcnb2^-/-^* mice. (G) Kaplan-Meier survival plot *Math1-Cre; SmoM2* with *Kcnb2^+/+^*, *Kcnb2^+/-^*, *Kcnb2^-/-^* alleles. (H) Representative images of P21 *Math1-Cre; SmoM2* and *Math1-Cre; SmoM2; Kcnb2^-/-^* mice displaying brain tumor-induced cranial bulging (left) and tumor burden (right). (I) Expression and quantification of *Kcnb2*, *Sox2*, and *Dcx* in a *Math1-Cre; SmoM2* tumor detected by RNAscope single-molecule in situ hybridization. (J-K) Nebulosa plot showing co-expression of *KCNB2* with *SOX2* and *DCX* in previously published human SHH MB single-cell RNA sequencing data^34, 35^.

### Kcnb2 is overexpressed in mouse SHH MB and regulates tumor progression

To determine the role of Kcnb2 in MB, we used the *Math1-Cre; SmoM2* genetic mouse model of SHH MB. SmoM2 is a constitutively active form of Smoothened, a key SHH signaling component^16^. Cre recombination in Math1-lineage cerebellar granule neuron progenitors, the SHH MB cell-of-origin, drives aberrant SHH signaling leading to aggressive brain tumors that recapitulate transcriptional and histopathological features of human SHH MB^17, 18^. Kcnb2 deficiency prolongs the survival of MB-bearing mice in an allelic dose-dependent manner, and Kcnb2 homozygous knockout results in a 50% improvement in survival accompanied by markedly reduced tumor burden compared to control MB-bearing mice (**Figure 3G-H**). All *Math1-Cre; SmoM2; Kcnb2^-/-^* mice developed MBs, indicating that Kcnb2 does not regulate tumor initiation.

Intratumoral heterogeneity in SHH MB is driven by hierarchically organized tumor cell types. Sox2^+^ or Olig2^+^ medulloblastoma-propagating cells (MPCs) are slow-cycling and resistant to anti-mitotic chemotherapy^32, 33^. Sox2^+^ MPCs give rise to Dcx^+^ transit-amplifying progenitor cells, which divide and differentiate into NeuN^+^ post-mitotic tumor cells^32^. Due to these properties, MPCs are a source of MB relapse and a Sox2^+^ gene signature portends poor prognosis in SHH MB patients^32^. We performed single molecule *in situ* hybridization and found that *Kcnb2* colocalizes with *Sox2^+^* as well as *Dcx^+^* cells in *Math1-Cre; SmoM2* tumors (**Figure 3I**). Furthermore, *KCNB2* was detected in two independent single cell RNA-seq datasets of human SHH MB containing a total of fourteen patient samples^34, 35^. Consistently, we observed co-expression of *KCNB2* with *SOX2* and *DCX* in human SHH MB (**Figure 3J-K**), raising the possibility that Kcnb2 regulates Sox2^+^ MPCs.

### Kcnb2 is required for tumor stage-dependent proliferative expansion of MPCs

We performed a series of pulse-chase experiments to investigate the label-retention properties of MPCs. The thymidine analogue BrdU is incorporated into DNA during S-phase and diluted upon serial cell divisions. To determine long-term MB cell cycle dynamics, BrdU was administered to tumor-bearing mice and analyzed following chase durations ranging from 1 to 14 days (**Figure S5A**). By quantifying BrdU label retention in distinct tumor populations, we found that Sox2^+^ MPCs are largely quiescent and label-retaining over the 14-day chase period (**Figure S5B** and **S5D**). Fast-cycling Dcx^+^ cells rapidly acquired and lost BrdU labelling (**Figure S5B** and **S5E**). Postmitotic NeuN^+^ cells are initially unlabeled, but NeuN^+^ ; BrdU^+^ cells slowly emerged concomitant with the decline of Dcx^+^ ; BrdU^+^ cells (**Figure S5B** and **S5F**), suggesting that NeuN^+^ cells inherited BrdU from differentiated Dcx^+^ cells. Therefore, our characterization of the quiescent, long-term label-retaining properties of Sox2^+^ cells, and lineage relationship between Dcx^+^ and NeuN^+^ cells in *Math1-Cre; SmoM2* MBs are consistent with findings from the *Ptch1^+/-^* model of SHH MB^32^. As *PTCH1* and *SMO* are the most commonly mutated genes in human SHH MB^36^, we conclude that MPC hierarchy is conserved in MB driven by distinct alterations in the SHH signaling pathway. Kcnb2 loss does not impact label retention dynamics of tumor cell types within the SHH MB hierarchy (**Figure S5C-F**), indicating that Kcnb2 knockout does not skew the Sox2^+^ hierarchy towards ectopic fates.

Next, we examined whether reduced tumor burden due to Kcnb2 loss is associated with alteration of Sox2^+^ MPCs. Since control MB-bearing mice display a median survival of 22 days, we examined the consequence of Kcnb2 knockout at early (P7) and late (P21) stages of MB tumorigenesis. In early-stage tumors, Kcnb2 knockout does not alter the number of MPCs (**Figure S4A**). In contrast, late-stage Kcnb2 knockout tumors display significantly fewer Sox2^+^ MPCs compared to control (**Figure 4A**). The reduced Sox2^+^ population was not attributable to apoptosis, as we did not detect differences in either apoptotic Sox2^+^ MPCs or tumor cells overall (**Figure S4E-G**). While the overall tumor mitotic index was comparable (**Figure S4C-D**), we observed stage-specific changes in the cycling status of Sox2^+^ MPCs (**Figure 4C-F**). In control tumors, the fraction of Ki67^+^ ; Sox2^+^ cycling MPCs decreased with tumor progression (P7 to P21; **Figure 4C-D**). Kcnb2 knockout diminishes MPC proliferation in early-stage (P7) MB (**Figure 4C-D**). At later stages (P21), both genotypes have comparably low levels MPC cycling (**Figure 4D**), consistent with a previous study that described the rapid proliferation of MPCs at SHH MB onset and transition towards quiescence in full-blown tumors^33^. These data demonstrate that Kcnb2 loss impairs the proliferative expansion of MPCs that occurs at early-stage MB.

**Figure 4.**
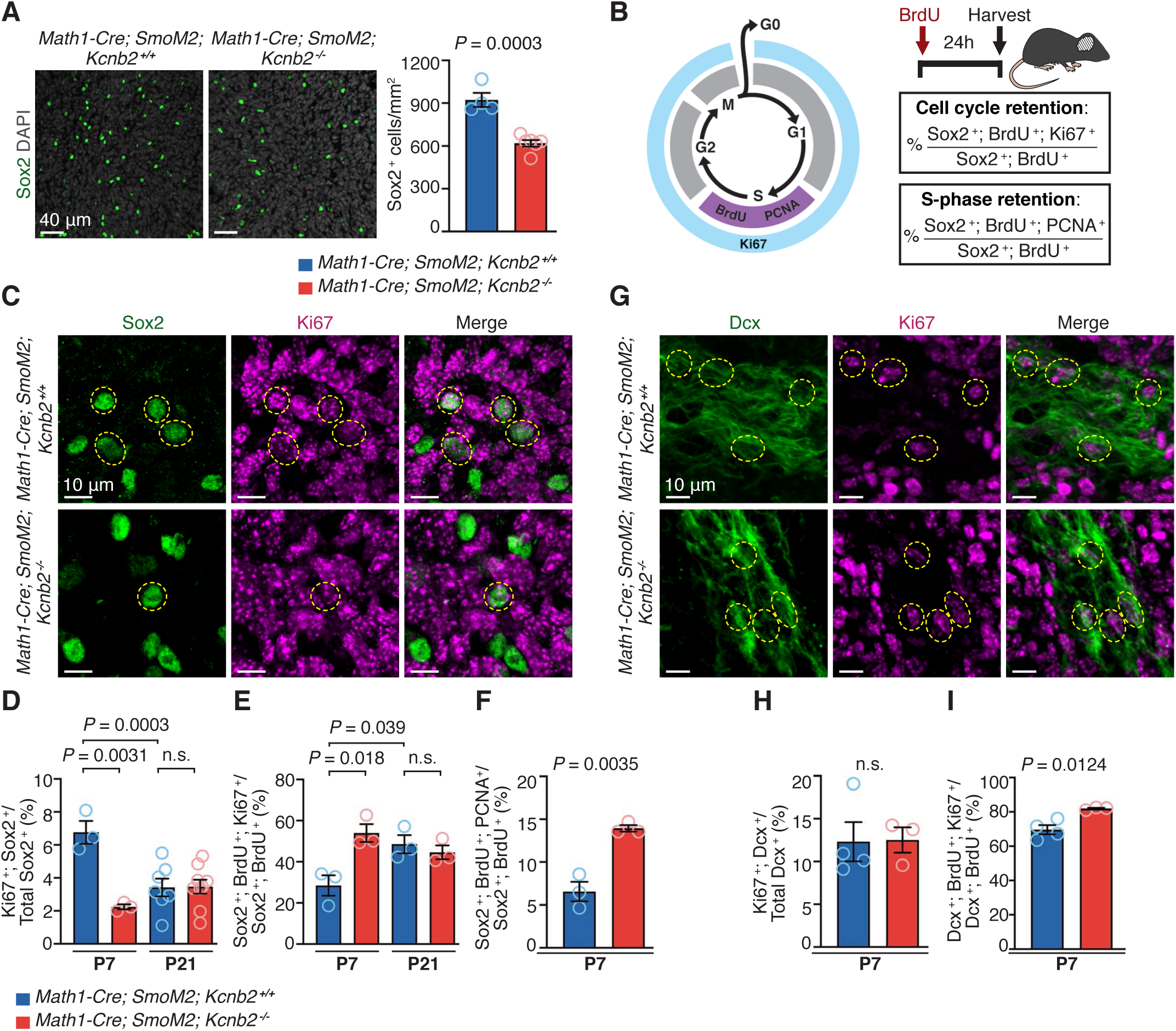
Kcnb2 regulates SHH MB-propagating cells. (A) Analysis of Sox2*^+^* cells in P21 MBs with or without Kcnb2 knockout. (B) Schematic of BrdU pulse experiment. (C) Representative images of Sox2 and Ki67 immunohistochemistry from P7 MBs. (D) Quantification of Ki67^+^ cycling proportion of Sox2^+^ MPCs at P7 and P21. (E) Quantification of Sox2^+^ cell cycle retention at P7 and P21. (F) Quantification of Sox2^+^ S-phase retention at P7. (G) Representative images of Dcx and Ki67 immunohistochemistry from P7 MBs. (H) Quantification of Ki67^+^ cycling proportion of Dcx^+^ MB cells at P7. (I) Quantification of Dcx^+^ cell cycle retention at P7.

To determine MPC cell cycle dynamics, MB-bearing mice were injected with BrdU and harvested 24 hours later (**Figure 4E**). By BrdU/Ki67 co-staining we found that P21 Sox2^+^ MPCs display significantly increased cell cycle retention compared to their P7 counterparts (**Figure 4E**). Consistent with Kcnb2 knockout not affecting the already-diminished proliferation of P21 MPCs (**Figure 4D**), cell cycle retention at P21 was not changed upon loss of Kcnb2 (**Figure 4E**). However, Kcnb2 knockout P7 MPCs display increased cell cycle retention compared to control (**Figure 4E**), indicative of an elongated cell cycle. BrdU/PCNA co-staining revealed that specifically, Kcnb2 knockout increases S-phase retention in P7 MPCs (**Figure 4F**). Collectively, these data show that in control MB, Sox2^+^ MPCs are more proliferative and undergo faster cell cycling during early tumor stages (P7) compared to later tumor stages (P21). Kcnb2 loss impairs this tumor stage-dependent proliferative expansion of the MPC pool by inducing S-phase retention and delaying cell cycle exit. Kcnb2 knockout does not impact the proliferation of transit-amplifying Dcx^+^ MB cells (**Figure 4G-H**), and leads to only a minor, and likely biologically insignificant increase in Dcx^+^ cell cycle retention (**Figure 4I**). Given that Kcnb2 knockout induces much stronger cell cycle defects in early stage Sox2^+^ MPCs, and their position atop the SHH MB hierarchy, we focused on determining the mechanism of Kcnb2 function in P7 Sox2^+^ MPCs.

### Kcnb2 regulates potassium homeostasis and mechanical properties of MPCs

To determine the mechanism by which Kcnb2 regulates MPCs, we isolated Sox2^+^ cells from MB^37^. Consistent with their *in vivo* defects, Kcnb2 knockout MPCs isolated from P7 mice display impaired *in vitro* growth (**Figure 5A**), as well as primary and secondary sphere-forming capacity (**Figure 5B**). We performed whole-cell patch clamp recordings to compare potassium channel activity in MPCs. Currents were elicited by voltage steps from -80 to 80 mV in 20-mV increments. Kcnb2 knockout MPCs display reduced potassium currents (**Figure 5C-D**). Administration of 4-Aminopyridine (4-AP), a blocker of voltage-gated potassium currents, to control MPCs phenocopies the knockout-induced sphere-forming defects (**Figure 5B**). These results indicate that Kcnb2-mediated potassium currents govern MPC proliferation in a cell-autonomous manner. Due to the higher intracellular potassium concentration, potassium channel opening causes potassium efflux along its electrochemical gradient. We postulate that Kcnb2 loss leads to elevated intracellular potassium, resulting in increased intracellular osmolarity to cause water influx and cell swelling. Consistent with this notion, Kcnb2 knockout MPCs display increased membrane capacitance (**Figure 5E**), which is proportional to the surface area of the plasma membrane^38^. Furthermore, Kcnb2 knockout MPCs develop larger cell size compared to control, as assessed by immunofluorescence for SmoM2-YFP (**Figure 5F**).

**Figure 5.**
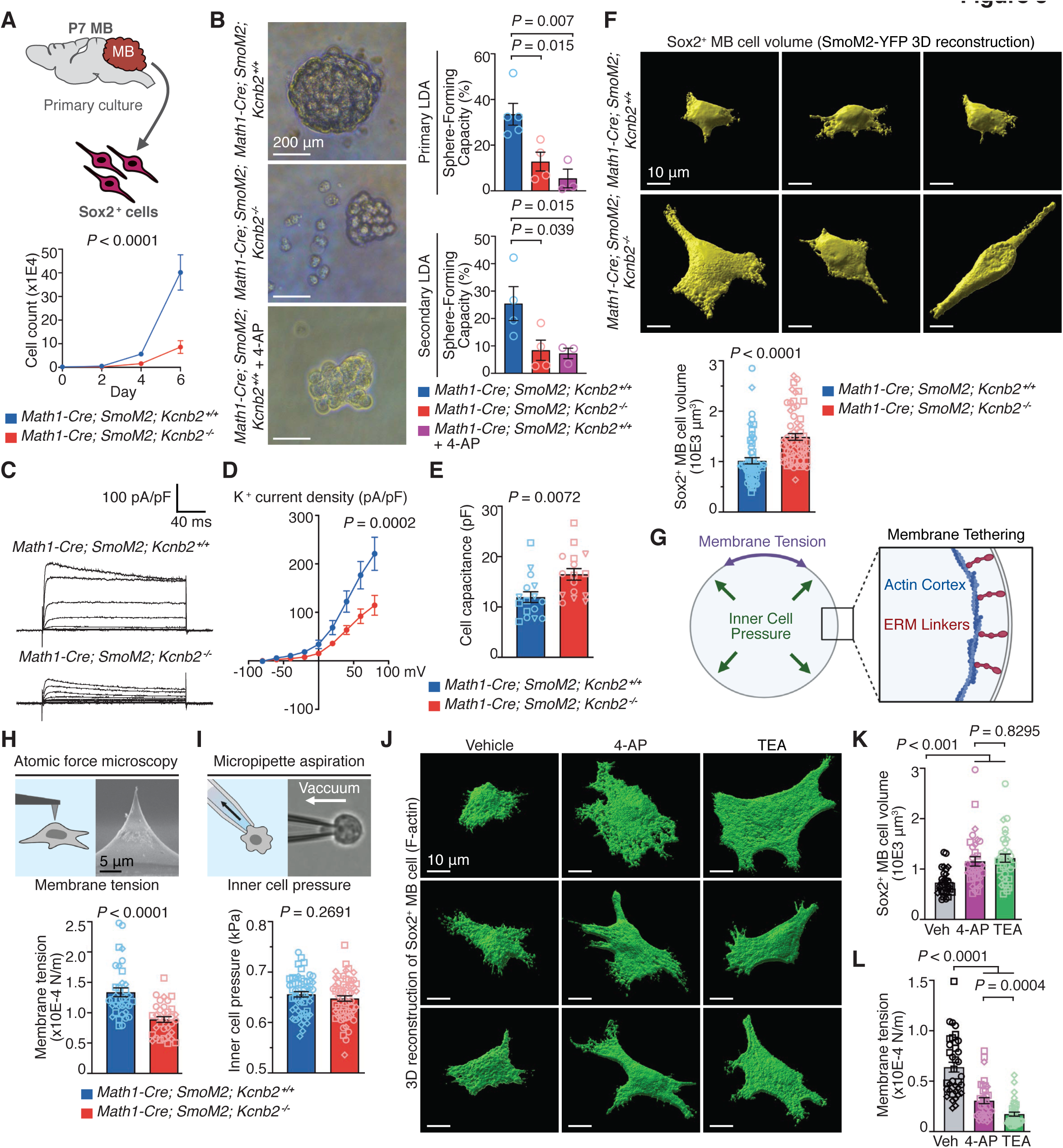
Kcnb2 regulates membrane tension of MB-propagating cells. (A) Top: Schematic for isolation and culture of Sox2^+^ MPCs. Bottom: 6-day cell counting assay comparing control and Kcnb2 knockout MPCs. (B) Primary and secondary *in vitro* sphere-forming limiting dilution analysis (LDA) comparing control and Kcnb2-knockout MPCs, and control MPCs treated with 2.5 mM 4-Aminopyridine (4-AP). (C) Representative current traces show total currents from whole-cell recordings of Sox2^+^ MPCs. Currents are elicited by voltage steps from -80 mV to 80 mV in 20-mV increments. (D) Current-voltage (I-V) curves of control and Kcnb2 knockout MPCs. (E) Cell capacitance of control and Kcnb2 knockout MPCs. (F) 3D-reconstruction and quantification of Sox2^+^ MPC volume as measured by SmoM2-YFP signal. (G) Schematic illustrating membrane tension, inner cell pressure, and membrane tethering. (H) Apparent membrane tension analysis of control and Kcnb2 knockout MPCs. (Top) Schematic and images shown for the probes used for atomic force microscopy experiments. (I) Inner cell pressure analysis of control and Kcnb2 knockout MPCs. (Top) Schematic and images show micropipette aspiration technique used. (J) Representative images of Sox2^+^ MPCs treated with 2.5 mM 4-aminopyridine (4-AP) and 5 mM tetraethylammonium (TEA). (K) Quantification of Sox2^+^ MPC volume as measured by α-Tubulin signal treated with 2.5 mM 4-aminopyridine (4-AP) and 5 mM tetraethylammonium (TEA). (L) Apparent membrane tension of Sox2^+^ MPCs treated with 2.5 mM 4-aminopyridine (4-AP) and 5 mM tetraethylammonium (TEA).

Given that Kcnb2 knockout impacts MPC size and plasma membrane surface area, we next investigated whether cell mechanics were impacted. Increased intracellular potassium due to Kcnb2 knockout and subsequent water influx may increase inner cell pressure, a force pushing outward against the plasma membrane. Plasma membrane constriction and expansion affect membrane tension (the in-plane force which counteracts surface expansion), a biophysical parameter which regulates cell fate specification^39, 40^. Therefore, we examined these two biomechanical parameters: membrane tension and inner cell pressure (**Figure 5G**). Using atomic force microscopy coupled with a sharp probe to induce small, local indentations, we found that Kcnb2 knockout reduces the plasma membrane tension of MPCs (**Figure 5H**). We determined inner cell pressure using micropipette aspiration, where a negative pressure is applied to MPCs. Inner cell pressure, which generates a rounding force to resist cell deformation, is inversely proportional to the amount of cell being aspirated into the micropipette. Kcnb2 knockout did not alter MPC inner cell pressure (**Figure 5I**), indicating that osmolarity-driven water influx and cell swelling may be compensated by modulating membrane tension. Broad inhibition of potassium currents with tetraethylammonium (TEA), a general potassium channel blocker, or inhibition of voltage-gated potassium currents with 4-AP increased cell size while reducing plasma membrane tension (**Figure 5J-L**). Taken together, these data show that Kcnb2-dependent potassium homeostasis regulates MPC proliferation, cell size, and plasma membrane tension.

### Kcnb2 regulates plasma membrane-actin cortex tethering of MPCs

Next, we investigated the mechanism by which Kcnb2 regulates plasma membrane tension. When cell size increases, plasma membrane remodels to ensure that the mechanically inflexible lipid bilayer does not rupture due to cell swelling. Plasma membrane-to-actin cortex tethering by ERM (Ezrin, Radixin, Moesin) proteins is a key contributing factor to cell surface mechanics, including membrane tension. Phosphorylation of ERM (pERM) induces an open conformation permissive to binding F-actin^41^. pERM membrane tethering sequesters membrane surface area to increase tension, whereas reduced pERM increases the amount of untethered membrane free to unfold in response to osmotic changes. We investigated if Kcnb2 regulates pERM membrane-actin tethering and membrane tension through changes in osmolarity, and tested whether manipulating extracellular osmolarity rescues changes in cell size and plasma membrane-actin tethering (**Figure 6A**). Under isotonic conditions, Kcnb2 knockout reduced pERM tethering of MPCs between the plasma membrane and the actin cytoskeleton, concomitant with increased cell size (**Figure 6B-C**) and decreased overall pERM protein (**Figure S6B**).

**Figure 6.**
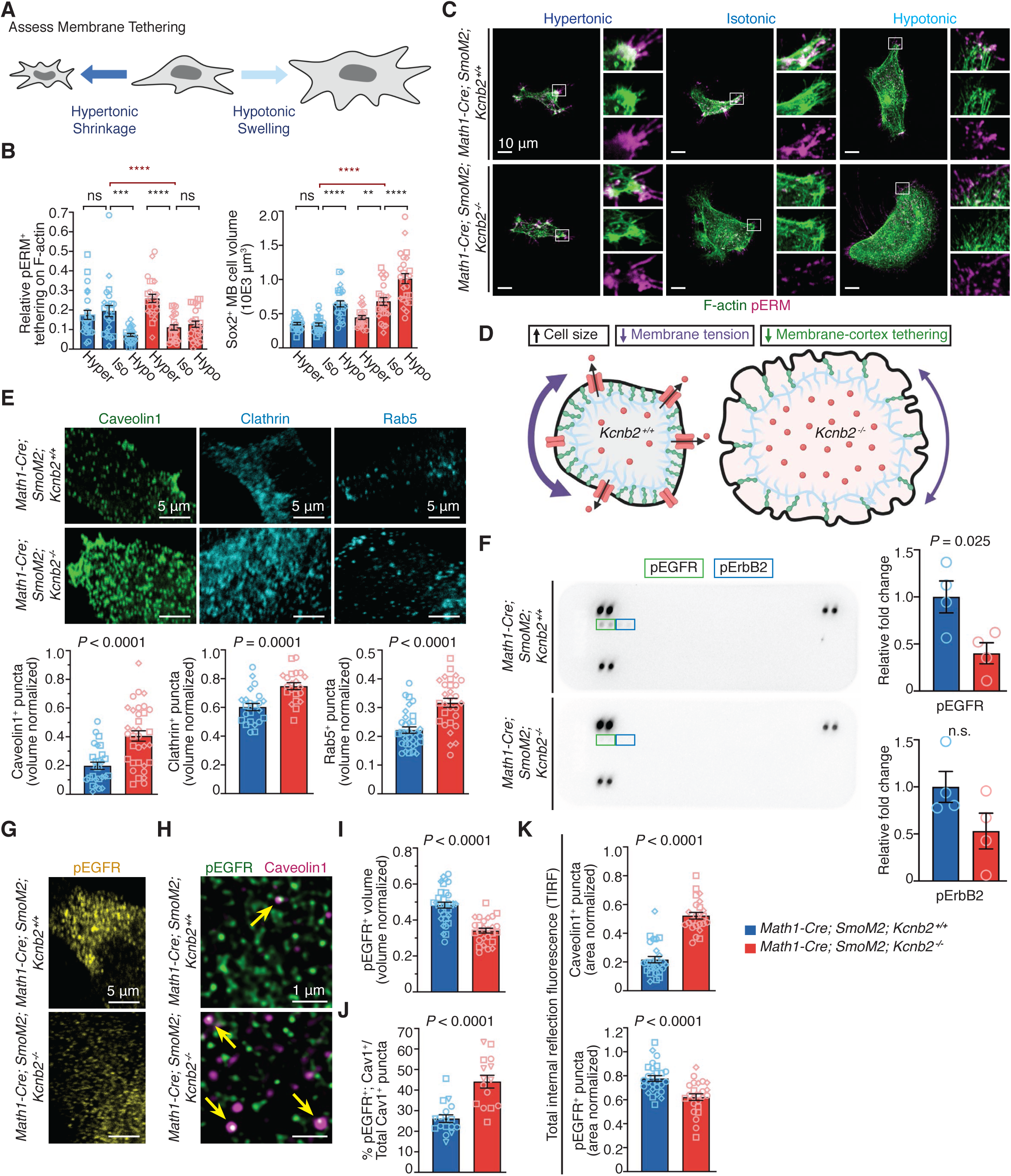
Kcnb2 regulates endocytosis of EGFR in MB-propagating cells. (A) Experimental schematics for manipulating osmolarity to assess membrane tethering. (B) Analysis of pERM membrane tethering and cell size in control and Kcnb2 knockout MPCs under hypertonic, isotonic, and hypotonic conditions. Quantifications show co-localized pERM^+^ ; F-actin^+^ signal normalized to total F-actin^+^ volume (left) and cell size as assessed by SmoM2-YFP volume (right). Hypertonic conditions were achieved by addition of 3.75% polyethylene glycol (PEG) 1500 for 24 hrs. Hypotonic conditions were achieved by addition of 30% de-ionized water for 24 hrs. (C) Representative immunocytochemistry for pERM and F-actin in wild-type and Kcnb2 knockout MPCs under hypertonic, isotonic, and hypertonic conditions. (D) Schematic representation of cell size and biomechanical changes upon Kcnb2 knockout. (E) Representative images and quantification of endocytic markers Caveolin-1, Clathrin, and Rab5 in Sox2^+^ MPCs. (F) Representative images and quantification of detectable pEGFR and pErbB2 signal from phospho-RTK array analysis of Sox2^+^ MPC lysates. (G) Representative images of phosphorylated EGFR (pEGFR) signal in Sox2^+^ MPCs. (H) Representative images of pEGFR^+^ and Caveolin-1^+^ colocalization in Sox2^+^ MPCs. (I) Quantification pEGFR signal in Sox2^+^ MPCs. (J) Quantification of pEGFR^+^ and Caveolin-1^+^ colocalization in Sox2^+^ MPCs. (K) Total internal reflection fluorescence (TIRF) microscopy and quantification of the membrane fraction of Caveolin-1 and pEGFR in Sox2^+^ MPCs.

Control MPCs in hypotonic media phenocopies Kcnb2 knockout, with cell swelling and reduced pERM tethering. However, hypertonic media failed to further shrink or increase membrane-actin tethering in control MPCs (**Figure 6B-C**). The enlarged cell size and reduced pERM tethering of Kcnb2 knockout MPCs is rescued by hypertonic conditions, which reduces cell size and increases membrane tethering. Conversely, while hypotonic conditions can further enlarge the already-swollen Kcnb2 knockout MPCs, this is not accompanied by further reduction in pERM tethering (**Figure 6B-C**). These data suggest that Kcnb2 maintains potassium and cell volume homeostasis to regulate MPC membrane tension through pERM-mediated membrane-actin tethering (**Figure 6D**). Membrane tension and pERM tethering rise as cells undergo mitosis to facilitate mitotic cell rounding and proper positioning of mitotic spindles by the tethered actin cortex^41, 42^. The proliferation defects and lengthened cell cycle observed in Kcnb2 knockout MPCs (**Figure 4C-F**) may be attributed to reduced MPC membrane tension and pERM tethering. The osmolarity manipulations suggest that osmotic regulation of cell size and membrane tension are regulated and coupled within a physiological range. Control Sox2^+^ MPCs are already quite small with very little cytoplasm, hindering further reduction in cell size. While hypotonic conditions can further increase the size of Kcnb2 knockout MPCs, coupling between cell size and pERM tethering breaks down, as further reduction of membrane tethering at large cell sizes may lead to membrane rupture.

### Kcnb2 regulates EGFR endocytosis through membrane tension

We sought to determine how membrane tension regulates MPC proliferation. Membrane tension serves as an energy barrier that gates biological processes involving membrane folding, such as endocytosis. A more flexible plasma membrane and reduced membrane tension facilitate endocytosis^40, 43, 44^. Endocytosis occurs mainly through clathrin- and caveolae-dependent pathways, where membrane domains to be internalized are encased by Clathrin and Caveolin-1, respectively. Kcnb2 knockout MPCs display increased Caveolin-1^+^ and Clathrin^+^ puncta, as well as increased Rab5^+^ puncta, a marker of early endosomes (**Figure 6E**). Endocytosis may regulate the signaling output of receptor tyrosine kinases (RTKs)^45^. We assessed RTK activation in MPC lysates through a phospho-RTK array, and only detected activation (phosphorylation) of epidermal growth factor receptor (EGFR) family members EGFR and ErbB2. Relative to control, Kcnb2 knockout MPCs exhibit reduced pEGFR, while pErbB2 signal was not significantly altered (**Figure 6F** and **S7A-C**). Brain tumor-propagating cells are traditionally maintained with epidermal growth factor (EGF) and fibroblast growth factor (FGF)^46^. Through a series of growth factor titrations, we found that MPC viability is crucially dependent on EGF but not FGF (**Figure S6A**), consistent with the absence of FGFR activation in the phospho-RTK array (**Figure 6F** and **S7A-C**). Our results suggest that MPC proliferation is primarily dependent on EGFR signaling.

Kcnb2 knockout MPCs display reduced amounts of activated phosphorylated EGFR (**Figures 6G**, **6I** and **S6B**), accompanied by increased colocalization of pEGFR^+^ and Caveolin-1^+^ puncta (**Figure 6H** and **6J**). Total internal reflection fluorescence (TIRF) microscopy, which specifically visualizes plasma membrane-localized proteins, further validated the increased Caveolin-1^+^ endocytic puncta and decreased levels of membrane-localized pEGFR in Kcnb2 knockout MPCs (**Figure 6K**). These data suggest that reduced membrane tension increases internalization of EGFRs through endocytosis in Kcnb2 knockout MPCs. We corroborated our results by immunohistochemistry of Sox2^+^ MPCs in early-stage (P7) *Math1-Cre; SmoM2* tumors. Consistently, Kcnb2 knockout MPCs display reduced pERM, increased Caveolin-1, and reduced pEGFR signal compared to control (**Figure 7A-B**), consistent with our *in vitro* observations.

**Figure 7.**
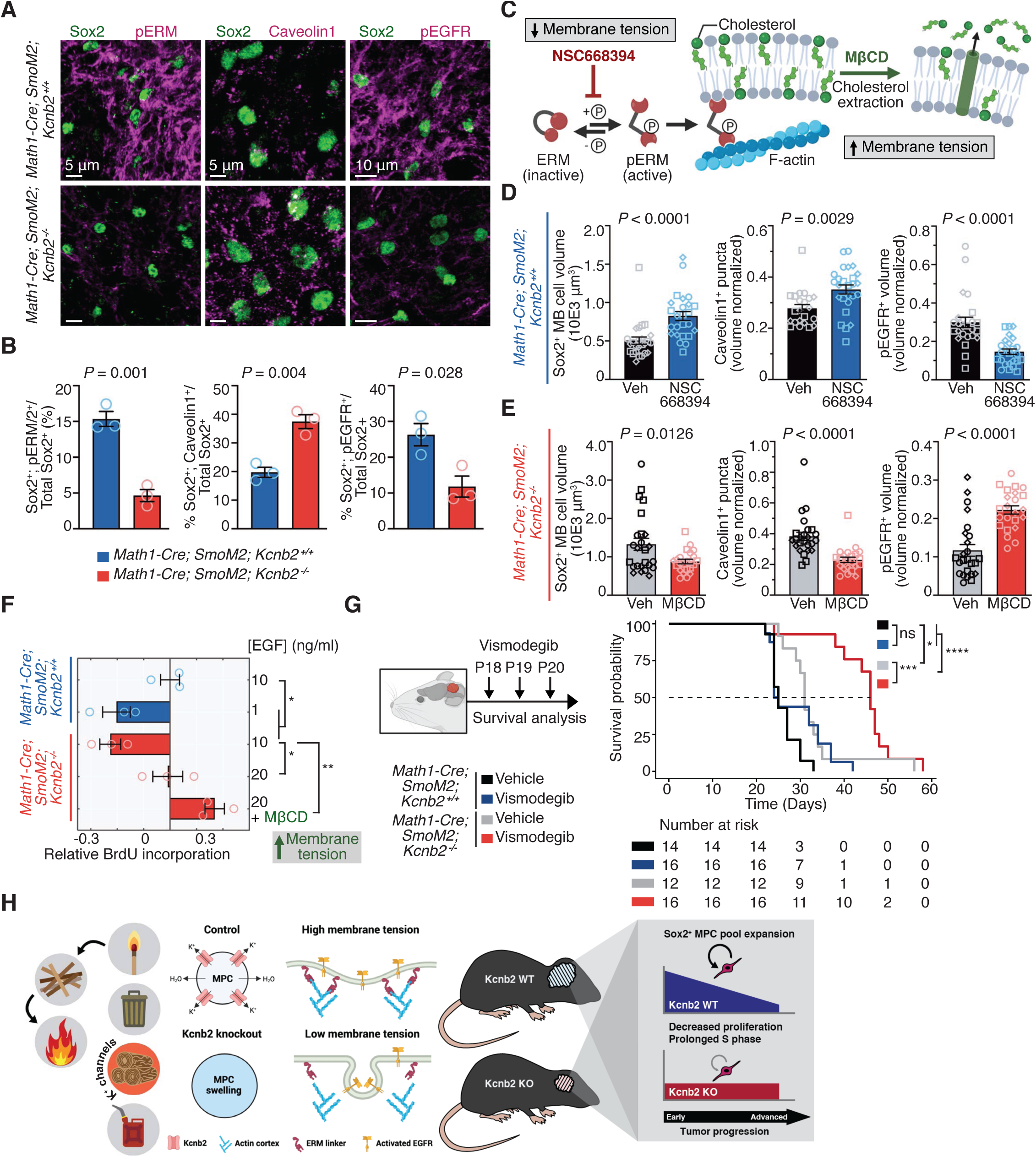
Ionic control of membrane tension and EGFR potentiate MB cell proliferation. (A-B) Representative immunohistochemistry (A) and quantification (B) of Sox2, pERM, Caveolin-1, and pEGFR, in P7 control and Kcnb2 knockout *Math1-Cre; SmoM2* MB. (C) Schematic of experiments to manipulate membrane tension. (Left) Control Sox2^+^ MPCs are treated with NSC668394, an inhibitor of ezrin phosphorylation, to reduce membrane tension. (Right) Kcnb2 knockout Sox2^+^ MPCs are treated with Methyl-β-cyclodextrin (MβCD), which depletes cholesterol from plasma membrane to increase membrane tension. (D) Quantification of SmoM2^+^ cell volume, Caveolin-1^+^ puncta, and pEGFR^+^ signal in control Sox2^+^ MPCs treated with 25 μM NSC668394 (NSC) for 24 hrs *in vitro*. (E) Quantification of SmoM2^+^ cell volume, Caveolin-1^+^ puncta, and pEGFR^+^ signal in Kcnb2 knockout Sox2^+^ MPCs treated with 1 mM MβCD for 24 hrs *in vitro*. (F) BrdU incorporation in Sox2^+^ MPCs after incubating for 3 hours. Results are expressed as fold change relative to control MPCs in standard culture conditions with 10 ng/mL EGF. (G) (Left) Schematic of vismodegib treatment regimen. (Right) Control and Kcnb2 knockout MB-bearing mice were treated on three consecutive days with vismodegib or vehicle and monitored for survival. (H) Physiological and molecular mechanism of action for Kcnb2 in MB.

We next manipulated membrane tension to establish causality between membrane tension, endocytosis, and EGFR signaling (**Figure 7C**). NSC668394, an inhibitor of Ezrin phosphorylation, reduces plasma membrane-to-actin cortex tethering and membrane tension^40, 47^. Methyl-β-cyclodextrin (MβCD) depletes cholesterol from the plasma membrane, reduces endocytosis, and increases membrane tension^40, 48, 49^. Treating control MPCs with NSC668394 increases cell size, elevates Caveolin-1 expression, and reduces pEGFR levels, mimicking the Kcnb2 knockout phenotype (**Figure 7D**). Conversely, treating Kncb2 knockout MPCs with MβCD reduces cell size, decreases Caveolin-1 expression, and increases pEGFR levels (**Figure 7E**). These data establish membrane tension as a regulator of endocytosis and EGFR signaling in MPCs.

### Ionic homeostasis and membrane tension govern EGFR signaling and proliferation in MB

Next, we asked whether manipulating the membrane tension–EGFR axis can rescue the proliferation defect of Kncb2 knockout MPCs. Relative to control, Kncb2 knockout MPCs have lower BrdU incorporation *in vitro* (**Figure 7F**). Decreased EGF concentration (from 10 ng/mL to 1 ng/mL) reduces BrdU incorporation in control MPCs to a similar extent as Kcnb2 knockout. Supplementation with EGF ligand rescues the BrdU incorporation rate of Kcnb2 knockout MPCs to control levels. Combination of EGF and MβCD further increases BrdU incorporation, suggesting that high membrane tension cooperates with EGF ligands to facilitate MPC proliferation (**Figure 7F**).

Finally, since we did not observe any developmental or physiological phenotypes in Kcnb2 knockout mice (**Figure 3A-F** and **S3A-F**), and Sox2^+^ MPCs are refractory to anti-SHH signaling therapies^32^, we determined whether Kcnb2 knockout synergizes with SHH pathway antagonism to treat SHH MB. Vismodegib, a small-molecule Shh signaling pathway antagonist, is approved for treating SHH signaling-driven basal cell carcinoma and undergoing clinical trials for recurrent SHH MB^50^. We administered three daily doses of vismodegib or vehicle to MB-bearing mice (**Figure 7G**). Vismodegib treatment alone did not improve the survival of control MB-bearing mice compared to vehicle treatment, consistent with a previous report^51^. In contrast, vismodegib treatment significantly prolonged the survival of Kcnb2 knockout MB-bearing mice. These data establish Kcnb2 as a target for the therapy-resistant MPCs, underscoring a synergism between Kcnb2 inhibition and tumor-debulking agents in treating SHH MB.

## DISCUSSION

A major obstacle in treating cancer is the ever-changing therapeutic vulnerabilities in space and time. Tumor evolution, which invariably occurs under the stress from tumor cells and their microenvironment, nullifies tumor dependencies on initiation mutations while conferring new dependencies that maintain the growing tumor. As a result, identifying mechanisms which maintain, rather than initiate, tumors are of tremendous clinical significance. Most cancer genomics studies are designed to identify tumor initiation events, leaving tumor maintenance underpinnings unexplored. In this study, we designed the Lazy Piggy insertional mutagenesis system to identify maintenance mechanisms of MB. In a genetic mouse model of MB, SB first mobilizes the hybrid transposon to disrupt gene expression. Subsequent PB activation depletes SB insertions, thereby restoring original gene function. Under these genomic manipulations, initiation, passenger, and progression insertions are removed with no effect on tumor survival. In contrast, removing maintenance insertions leads to clonal collapse. Therefore, Lazy Piggy functional genomics filters out initiation, passenger, and progression genes, while revealing the maintenance mechanisms. This represents the first *in vivo* functional genomics tool designed to uncover tumor maintenance events.

Using this novel insertional mutagenesis approach, we identify potassium homeostasis as a critical dependency of MB. *KCNB2* is the most consistently upregulated potassium channel across all four MB subgroups. While Kcnb2 is non-essential for normal mouse development, Kcnb2 knockout markedly hinders MB growth and prolongs survival of tumor-bearing mice. Kcnb2 expression is enriched in the therapy-resistant Sox2^+^ MPCs, and Kcnb2 loss diminishes the MPC pool by impairing its tumor stage-dependent proliferative expansion. Mechanistically, Kcnb2 maintains MPC cell size and governs the mechanical properties of plasma membrane, which in turn gates endocytosis-regulated EGFR signaling (**Figure 7H**). Plasma membrane tension is a tightly controlled biomechanical property which governs cell behaviors such as migration and division^52^. In embryonic stem cells, reduction of membrane tension is an essential step in differentiation and pluripotency exit^39, 40^. Here, our study establishes Kcnb2 as a vulnerability in MB maintenance and provides the first evidence that ionic homeostasis-gated membrane tension is critical for tumor-propagating cell proliferation and tumor progression. Furthermore, we show that targeting Kcnb2 synergizes with tumor debulking chemotherapies to impair MB growth.

Osmotic stress is a physiologically relevant force which cells must adapt and respond to. The relatively inextensible plasma membrane can only sustain 3-4% stretch before membrane rupture^53^, and thus membrane tension must be maintained at sub-lytic levels to prevent cell lysis. In this study, we found that Kcnb2 knockout in MPCs leads to hypotonic cell swelling (**Figure 5F**) and reduced membrane tension (**Figure 5H**). This contrasts with previous studies reporting that membrane tension spikes within seconds of acute hypotonic cell swelling, followed by recovery of both volume and tension on a timescale of minutes to hours^54, 55^. Excess membrane is stored as cell surface reservoirs of protrusions and invaginations, maintained by cytoskeleton-membrane tethering^56^. The initial transient tension increase reflects an inability to rapidly provide excess membrane and counterbalance the increased cell volume^57^. Cells then responds to hypotonic swelling by cytoskeletal reorganization to release tethered membrane surface area and drive membrane unfolding^55^. Finally, recovery from hypotonic swelling is mediated by ion channels and transporters which efflux osmolytes from cells to reduce volume. Being the major intracellular cation class, potassium has a dominant influence in establishing electrochemical equilibrium. Global and constitutive Kcnb2 loss from MPCs leads to reduced outward potassium currents (**Figure 5C-D**) and increased cell size (**Figure 5F**), indicating that compensatory mechanisms are not sufficient to rescue the perturbed potassium homeostasis. Since Kcnb2 knockout MPCs remain larger than control, our data suggest that the membrane reservoirs remain depleted, cytoskeleton-membrane tethering and membrane refolding cannot occur, thus resulting in decreased membrane tension.

Previous studies of extrinsic osmotic shock and membrane tension reflect acute and transient cellular adaptations to hypotonic swelling in cells with intact volume regulatory machinery^54, 55, 57^. Our study differs in three critical aspects. First, we induced osmotic shock in a cell-autonomous manner by knocking out the endogenously overexpressed potassium channel Kcnb2. Second, we describe the chronic effects of dysregulated potassium homeostasis on membrane tension in cells unable to re-establish their original ionic balance and cell volume. Finally, intrinsic differences between the various cell types studied must be taken into consideration. The prior studies used *Lymnaea* molluscan neurons, Madin Darby canine kidney (MDCK II) cells, and HeLa Kyoto cancer cells respectively^54, 55, 57^, while our mechano-osmotic experiments were performed in murine brain tumor cells. While these studies all provide critical insights into the interplay between osmoregulation and mechanobiology, the cell-intrinsic differences preclude comparisons across species.

While specific ion channels are overexpressed in cancer, their expression in non-malignant tissues is tightly regulated, providing a therapeutic window to target ion channels in cancer^26^. While calcium is a well-established second messenger that engages intracellular signaling, how other cations such as potassium influence mechano-chemical signaling was unclear. Our study reveals how potassium homeostasis regulates plasma membrane-actin cortex tethering and cell surface mechanics. Decreased membrane tension enhances endocytosis, which enhances internalization of membrane-localized signaling molecules such as EGFR and attenuates the signaling output. Since membrane tension reduction leads to a more ‘pliable’ plasma membrane, we emphasize the possibility that the membrane tension drop we observe in Kcnb2 knockout BTICs creates a permissive environment for other processes dependent on plasma membrane remodeling, such as pinocytosis and exocytosis. While we implicate EGFR signaling as a functionally relevant consequence of Kcnb2 knockout-induced endocytosis, EGFR is likely not the sole membrane protein affected by alterations in plasma membrane tension.

Lastly, our study reveals that osmolarity influences plasma membrane tension. MB, as many other tumor types, develops extensive intratumoral heterogeneity. Our study raises the possibility that microenvironment-dependent differences in extracellular osmolarity may dictate membrane tension and tumor cell behavior. The extracellular osmotic environment in MB can differ depending on whether tumor cells are exposed to cerebrospinal fluid near the brain ventricles, plasma near leaky blood vessels, or interstitial fluid in the densely packed tumor core. An intriguing question is whether distinct membrane tension levels arise in tumor cells situated in anatomically different tumor regions, and whether potassium channels integrate osmotic and mechanical heterogeneity to regulate functional plasticity in stemness, lineage bias, and therapy resistance. Potassium homeostasis-governed biophysical properties of cancer represent therapeutic avenues to halt tumor progression.

## LIMITATIONS OF THE STUDY

Functional cancer genomics enables the identification of genes which may be missed by expression analysis. However, *in vivo* genomic approaches may not distinguish between tumor-essential and pan-essential genes, with the latter resulting in a limited therapeutic index. Conversely, differential expression analysis may yield tumor-specific targets that require functional validation. We partially circumvented this limitation by focusing on enriched pathways to reveal convergence on the potassium channel family as mediators of MB maintenance, and through analysis of MB patient expression data identify KCNB2 as the most consistently upregulated potassium channel in MB. A limitation of our study is the small sample size of Lazy Piggy tumors, which may have contributed to the discordant identification of KCNB2 from patient expression analysis but not the Lazy Piggy screen. Further studies using the Lazy Piggy model will require a larger cohort to increase genomic coverage of transposon mobilization and remobilization to discover new tumor maintenance drivers.

## ACKNOWLEDGEMENTS

This work is supported by Sontag Foundation Distinguished Scientist Award, Early Researcher Award, Canadian Cancer Society Challenge Grant, Cancer Research Society Operating Grant, Natural Sciences and Engineering Research Council Discovery Grant, American Brain Tumor Association Discovery Grant, Ontario Institute for Cancer Research Translational Research Initiative, Canadian Institute of Health Research Project Grants, b.r.a.i.n.child, and Meagan’s HUG to X.H. and National Institutes of Health grants CA148699 and CA159859, Stand Up To Cancer (SU2C) St. Baldrick’s Pediatric Dream Team translational research grant (SU2C-AACR-DT1113) and SU2C Canada Cancer Stem Cell Dream Team research funding (SU2C-AACR-DT-19-15) provided by the Government of Canada through Genome Canada and the Canadian Institutes of Health Research, with supplementary support from the Ontario Institute for Cancer Research through funding provided by the Government of Ontario to M.D.T. SU2C is a program of the Entertainment Industry Foundation administered by American Association for Cancer Research. M.D.T. is also supported by Pediatric Brain Tumor Foundation, Terry Fox Research Institute, Canadian Institutes of Health Research, Cure Search Foundation, b.r.a.i.n.child, Meagan’s Walk, SWIFTY Foundation, Brain Tumour Charity, Genome Canada, Genome BC, Genome Quebec, Ontario Research Fund, Worldwide Cancer Research, V-Foundation for Cancer Research, and Ontario Institute for Cancer Research through funding provided by the Government of Ontario. Additionally, M.D.T. is supported by Canadian Cancer Society Research Institute Impact grant, Cancer Research UK Brain Tumour Award, Garron Family Chair in Childhood Cancer Research at The Hospital for Sick Children, and University of Toronto. J.J.F. is supported by Restracomp Scholarship from The Hospital for Sick Children. X.W. is supported by Lap-Chee Tsui Fellowship from The Hospital for Sick Children and Dunn with Cancer Research Fellowship from Brain Tumour Foundation of Canada. A.W.E. is supported by Restracomp Fellowship from The Hospital for Sick Children. X.C. is supported by Restracomp Fellowship from The Hospital for Sick Children. S.B. is supported by International Postdoc Grant from Swedish Research Council. N.A. is supported by Childhood Cancer Research Postdoctoral Fellow Grant from Rally Foundation and Restracomp Fellowship from The Hospital for Sick Children. We thank Paul Paroutis and Kimberly Lau at SickKids Imaging Facility for help with confocal imaging and image analysis. Schematics were created using BioRender. X.H. thanks the love and support from Lucas Huang and Liam Huang. X.H. is a Catalyst Scholar at The Hospital for Sick Children and Canada Research Chair in Cancer Biophysics.

## AUTHOR CONTRIBUTIONS

J.J.F. performed all mouse work and wet laboratory experiments; Xin W. generated and performed validation experiments of the LP mouse; P.S. and A.W.E. were responsible for LP bioinformatic analysis; Xian W. performed membrane tension measurements; G.S. performed micropipette aspiration experiments; X.C. performed electrophysiology experiments and assisted with mouse work; Y.X., S.W., H.F., and A.S.M. assisted with bioinformatic analysis; S.B., M.A.F., R.J.P., and N.A. assisted with mouse work; all other authors contributed to methodology development and reagents. J.J.F. and A.W.E. wrote the manuscript with input from all other authors; X.H. and M.D.T. conceived and supervised the project.

## DECLARATION OF INTEREST

The authors declare no competing financial interests.

## FIGURE LEGENDS

**Supplemental figure 1.**
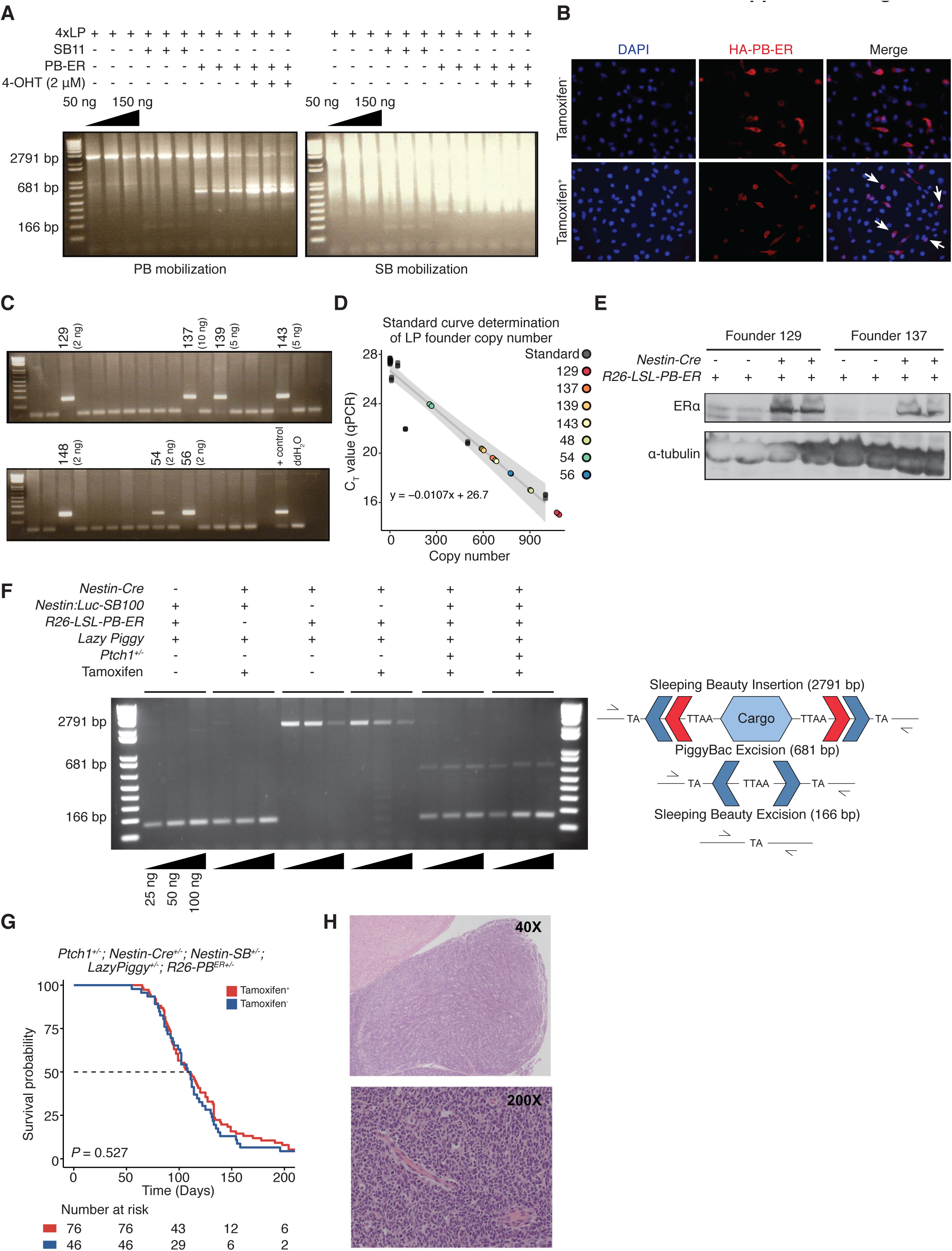
Generation and validation of transgenic mice harboring Lazy Piggy transposon. (A) PCR amplification of LP concatemer region in transfected 293T cells. Combinations performed in triplicate with 50ng, 100ng, and 150ng input DNA each. (B) Immunofluorescence of 293T cells transfected with LP transposon and PB-ER transposase demonstrated nuclear localization of PB transposase following tamoxifen administration. (C) Microinjection of 4-unit concatemer of the LP transposon generated 7 founder mice. (D) qPCR determination of LP concatemer copy number in founder mice. (E) Western blot analysis of cerebellar ERα expression in *R26-LSL-PB-ER^T2+/-^* mice when crossed with *Nestin-Cre^+/-^* mice. (F) PCR amplification of LP concatemer region demonstrates SB and PB mobilization with tamoxifen treatment in *Ptch1^+/-^; Nestin:Luc-SB100^+/-^; Lazy Piggy^+/-^; Nestin-Cre^+/-^; R26-LSL-PB-ER^T2+/-^* mice. Combinations performed in triplicate with 50ng, 100ng, and 150ng input DNA each. (G) Tamoxifen treatment was not associated with overall survival in *Ptch1^+/-^; Nestin:Luc-SB100^+/-^; Lazy Piggy^+/-^; Nestin-Cre^+/-^; R26-LSL-PB-ER^T2+/-^* mice. (H) Tumors in *Ptch1^+/-^; Nestin:Luc-SB100^+/-^; Lazy Piggy^+/-^; Nestin-Cre^+/-^; R26-LSL-PB-ER^T2+/-^* mice histologically resembled human medulloblastoma (H&E staining).

**Supplemental figure 2.**
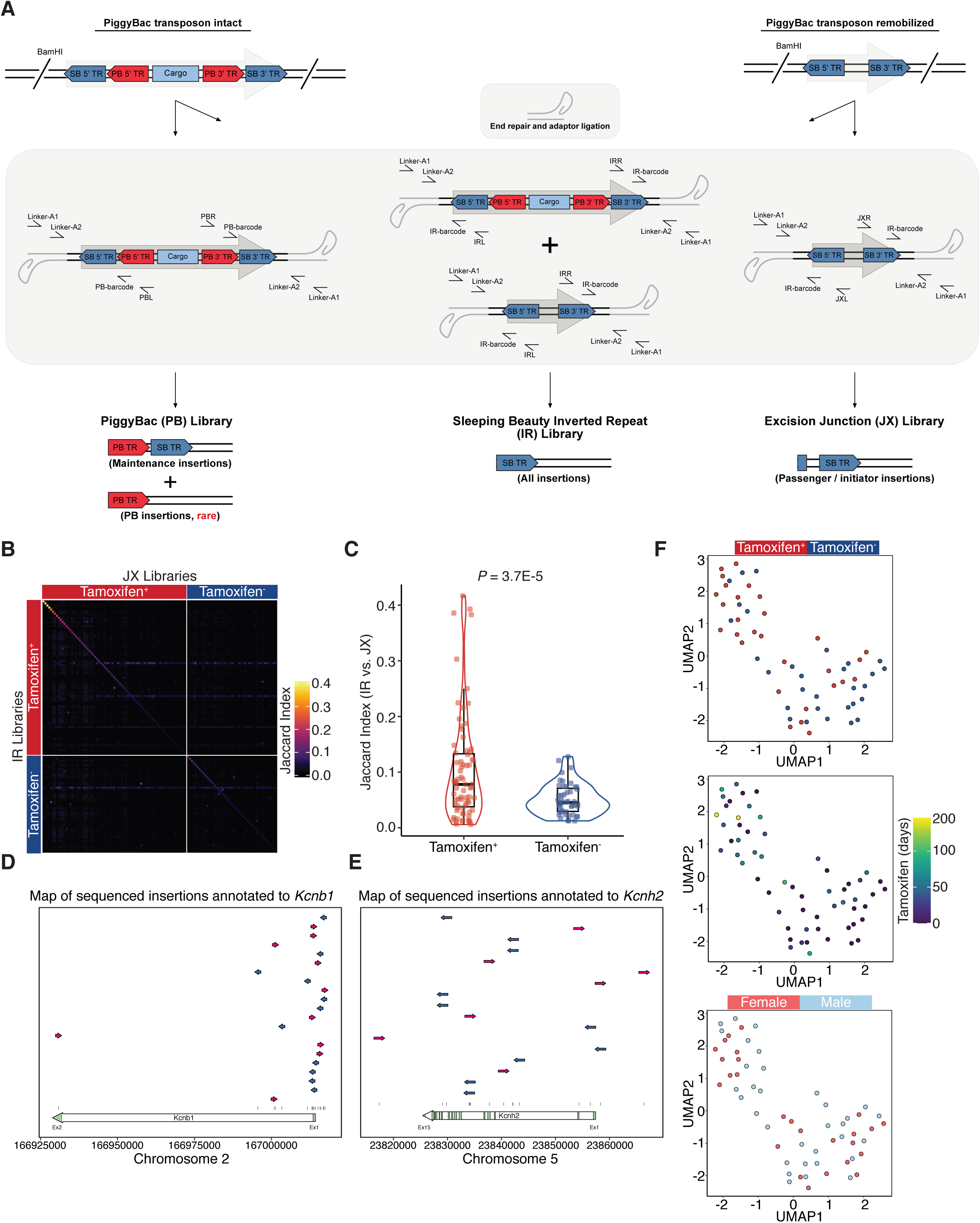
Detection and analysis of Lazy Piggy transposon remobilization. (A) Cartoon illustrating library preparation workflows for SB Inverted Repeat (IR), piggyBac (PB), and Excision Junction (JX) libraries. (B) Heatmap of Jaccard score similarity matrix for clonal IR and JX library insertions in sequenced mouse tumors. (C) Tamoxifen-treated tumors show a higher proportion of IR/JX overlap than untreated tumors. (D) Map of all sequenced insertions annotated to *Kcnb1* suggests gain-of-function given biases toward intronic, 5’, and sense-oriented insertions. (E) Map of all sequenced insertions annotated to *Kcnh2* suggests loss-of-function given the span of insertions across the entire coding sequence of *Kcnh2* which would yield a truncated and non-functional channel. (F) Dimensionality reduction by uniform manifold approximation and projection (UMAP) of DESeq2-normalized RNA-seq counts from tamoxifen-treated and untreated tumors.

**Supplemental figure 3.**
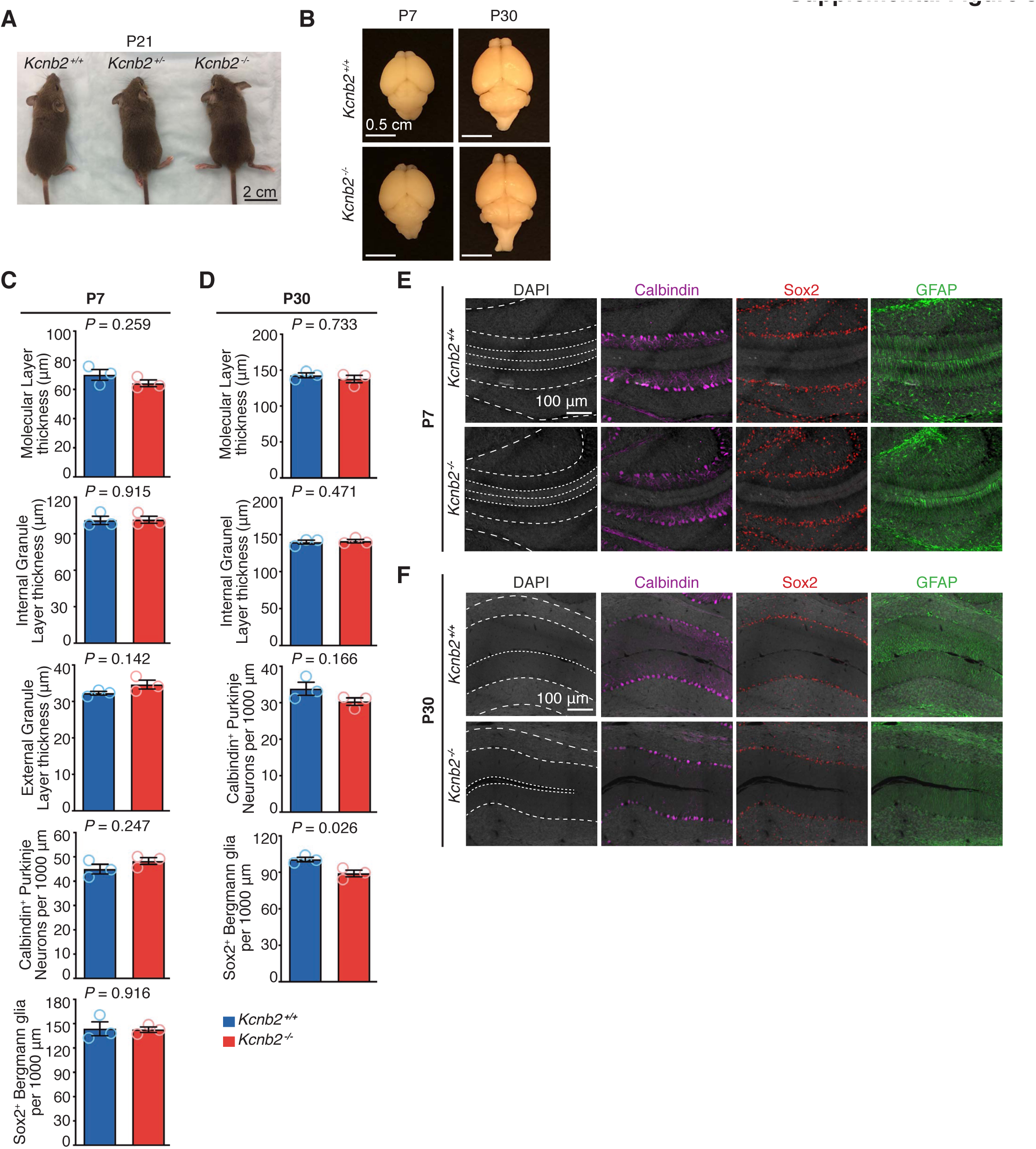
Analysis of Kcnb2 function in mouse cerebellar development. (A) Representative images P21 *Kcnb2^+/+^*, *Kcnb2^+/-^*, and *Kcnb2^-/-^* littermate mice. (B) Representative images of P7 and P30 brains of *Kcnb2^+/+^* and *Kcnb2^-/-^* mice. (C) Quantification of molecular layer, internal granule layer, external granule layer thickness, Calbindin^+^ Purkinje neuron and Sox2^+^ Bergmann glia populations of P7 *Kcnb2^+/+^* and *Kcnb2^-/-^* mice. (D) Quantification of molecular layer, internal granule layer thickness, Calbindin^+^ Purkinje neuron and Sox2^+^ Bergmann glia populations of P30 *Kcnb2^+/+^* and *Kcnb2^-/-^* mice. (E) Representative immunohistochemistry images from cerebella of P7 *Kcnb2^+/+^* and *Kcnb2^-/-^* mice. (F) Representative immunohistochemistry images from cerebella of P30 *Kcnb2^+/+^* and *Kcnb2^-/-^* mice.

**Supplemental figure 4.**
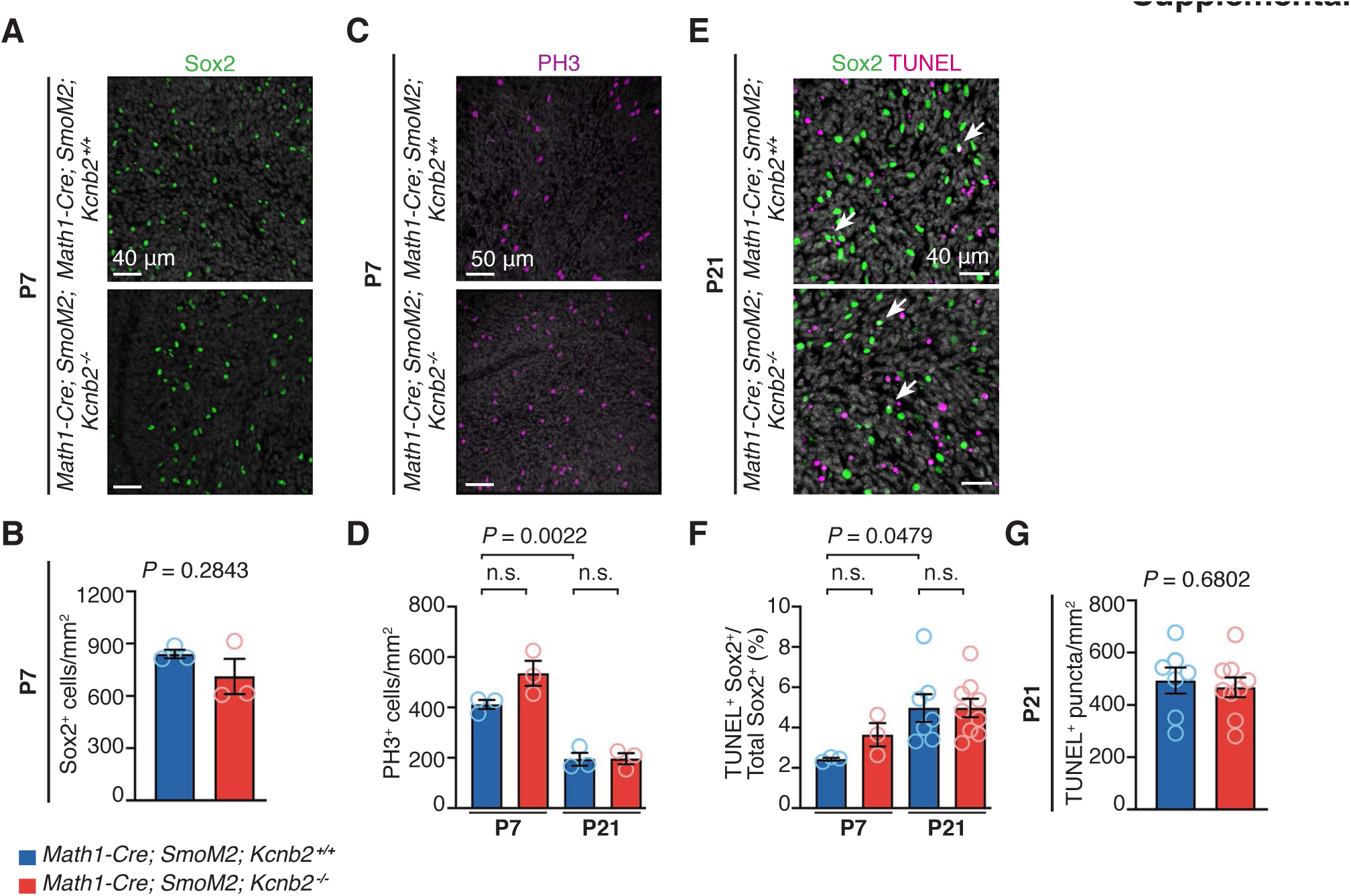
Phenotypic analysis of control and Kcnb2 knockout MB. (A-B) Quantification of Sox2^+^ cells in MB of P7 *Math1-Cre; SmoM2* mice with and without Kcnb2. (C-D) Quantification of phosphor-Histone 3^+^ (pHis3^+^) cells in MB of P7 *Math1-Cre; SmoM2* mice with and without Kcnb2. (E) Representative immunohistochemistry of TUNEL^+^ puncta and Sox2^+^ cells in MB of P21 *Math1-Cre; SmoM2* mice with and without Kcnb2. (F) Quantification of TUNEL^+^; Sox2^+^ cells in P7 and P21 MB of *Math1-Cre; SmoM2* mice with and without Kcnb2. (G) Quantification of TUNEL^+^ tumor cells in P21 MB of *Math1-Cre; SmoM2* mice with and without Kcnb2.

**Supplemental figure 5.**
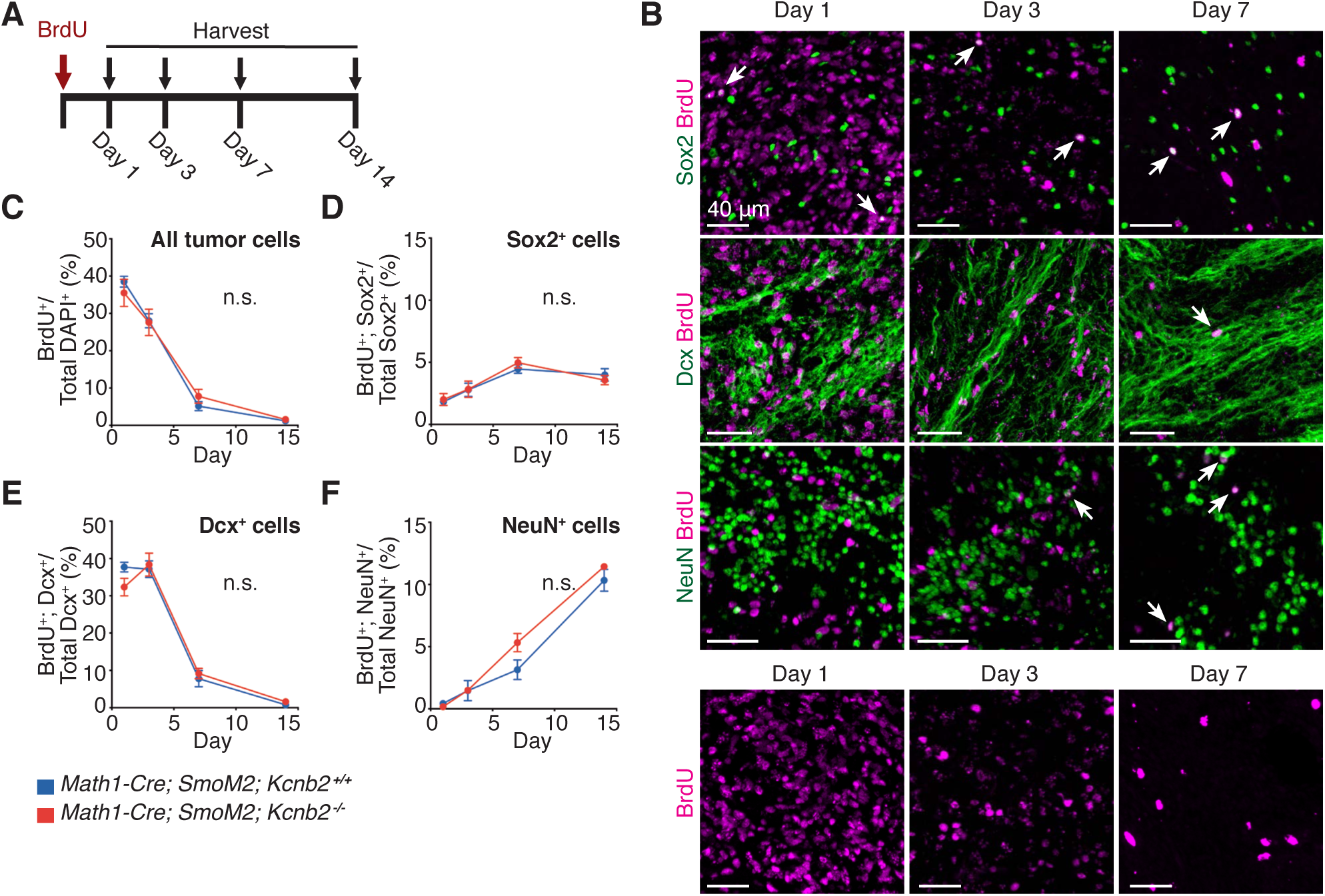
BrdU label retention analysis of MB. (A) Experimental design of BrdU label retention analysis. *Math1-Cre; SmoM2* and *Math1-Cre; SmoM2; Kcnb2^-/-^* mice were injected with a single dose of BrdU and sacrificed at the indicated timepoints (*n* = 3-4 mice per group). (B) Representative images of BrdU label retention in Sox2^+^, Dcx^+^, and NeuN^+^ cells at 1, 3, and 7 days after BrdU injection in MB of *Math1-Cre; SmoM2* mice. (C-F) Percentage of BrdU retention in all tumor cells (C), Sox2^+^ cells (D), Dcx^+^ cells (E), and NeuN^+^ cells (F).

**Supplemental figure 6.**
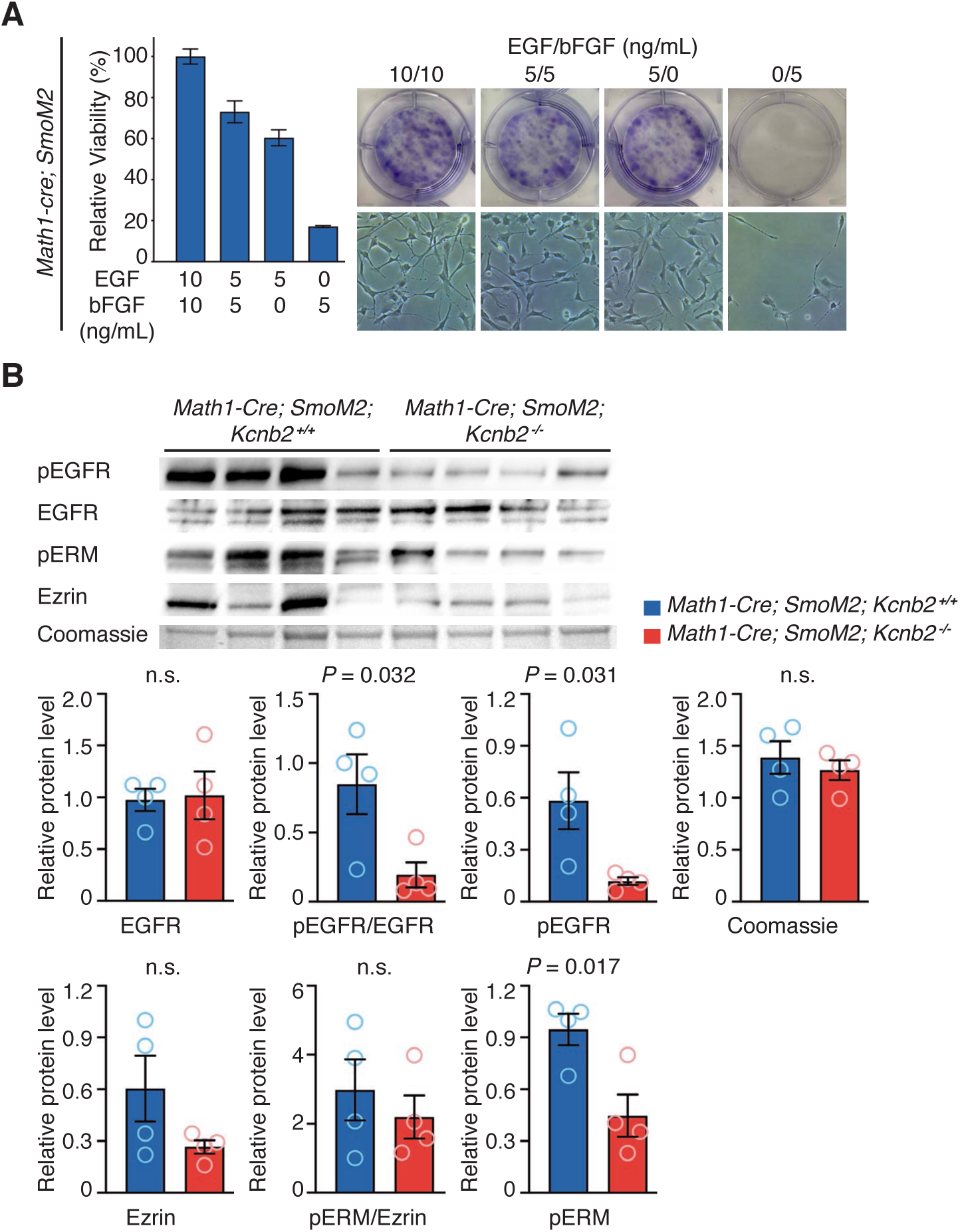
Ligand titration assay and western blot in Sox2^+^ MPCs *in vitro*. (A) Sox2^+^ MPCs were cultured in media with indicated concentrations of EGF and bFGF and assessed for relative viability by MTS assay (left) and crystal violet staining (right). (B) Western blot and quantification of four pairs of control and Kcnb2 knockout MPCs for pEGFR, EGFR, pERM, Ezrin, and Coomassie blue dye. All quantifications were normalized to the total Coomassie blue signal per lane.

**Supplemental figure 7.**
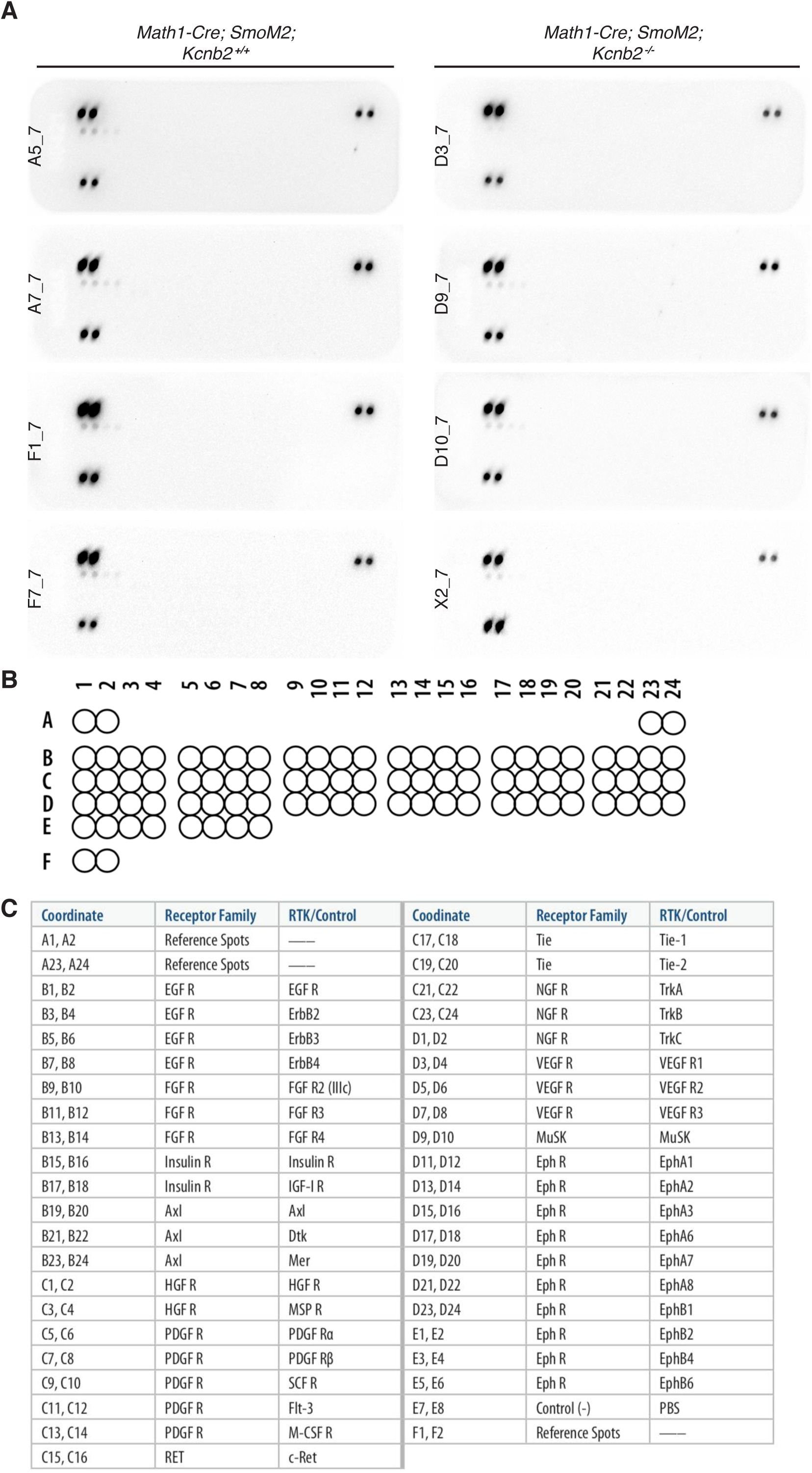
Phospho-RTK array of Sox2^+^ MPCs *in vitro*. (A) Phospho-RTK array of control and Kcnb2 knockout Sox2^+^ MPCs. Each array was performed on a separate biological replicate. (B) Phospho-RTK array map. (C) Phospho-RTK array legend.

## METHODS

### Lazy Piggy plasmid construction

The Lazy Piggy transposon was custom synthesized and is shown in Figure 1B. Mammalian-optimized PB minimum ITRs, as previously published^59^, were introduced to the T2/Onc2 SB transposon, nested between SB ITRs. T2/Onc2, as previously published^2^, contained a murine stem cell virus long terminal repeat (MSCV) 5’ LTR, a splice donor (SD) from exon 1 of the mouse Foxf2 gene, one splice acceptor (SA) from exon 2 of the mouse engrailed-2 (En2) gene and another from the carp β-actin gene, followed by a bidirectional SV40 poly(A) sequence. Restriction sites AcII and ClaI were introduced flanking the transposon sequence and a 4-copy concatemer was cloned into the pUC19 vector, hence termed “pLazyPiggy.” Detailed cloning methods available upon request.

### *In vitro* validation of the Lazy Piggy system

HEK293T cells were cultured in DMEM (GIBCO) supplemented with 10% fetal bovine serum at 37°C and 5% CO_2_. 5 × 10^5^ cells were seeded into each well of a 6-well plate 1 day prior to transfection. For each well, 2µg circular plasmids pCMV-HA-mPB-ERT2^59^, pLazyPiggy, and pCMV-SB11^11^ were transfected using Lipofectamine LTX as per manufacturer’s protocol (Invitrogen). Cells were then incubated with 2 μM 4-hydroxytamoxifen (4-OHT) for 24 hours. Cells were then collected and gDNA were extracted for PCR excision assays. Primers for amplifying LP-transposon mobilization were based on the cloning vector sequences adjacent to the inverted repeats/direct repeats (left) (IRDRL) and inverted repeats/direct repeats (right) (IRDRR) of the LP transposon, 5’-CGTTCACGACGTTGTAAAACGACG-3’ and 5’-CGATAATTAACCCTCACTAAAGGG-3’, respectively. The input represents genomic DNA with LP intact transposon (2791bp), post-SB mobilization (166bp) and post-PB re-mobilization (681bp) in the presence of 4-OHT. Detailed PCR protocol methods are available upon request.

### Novel mouse line generation

*Nestin-Cre* transgenic mice obtained from Jackson Laboratories (JAX stock #003771). To generate *Nestin:Luc-SB100* transgenic mice, SB100 cDNA was excised from the vector pCMV-SB100^60^, luciferase-tagged, then inserted into nes1852tk/lacZ plasmid, which carried the Nestin second intronic enhancer that has previously been shown to drive transgene expression in CNS stem and progenitor cells^61^. *Rosa26-LSL-mPB-L3-ERT2* transgenic mice were obtained from Dr. Allan Bradley^62^. Lazy Piggy transgenic mice were produced by pronuclear microinjection into zygotes by the Transgenic Core at The Centre for Phenogenomics (TCP)^63^. High-percentage male chimeras were crossed to C57/BL6 females. Germline transmission was confirmed by genotyping F1 offspring tail-clipped DNA. Primers used were: LP Fwd 5’-CGATAAAACACATGCGTC -3’, LP Rev 5’-CTCCAAGCGGCGACTGAG -3’.

Lazy Piggy founder copy number was determined through qPCR as previously described^64^. Briefly, a linear equation was modelled of CT values against copy number from 9 known LP transposon plasmid standards in triplicate, then CT values from triplicate founder samples were input to this model and resulting predicted copy numbers were averaged. Two founders (#129 and #137) were chosen for having “high” and “medium” LP copy numbers, respectively, on separate donor chromosomes.

### Western blot analysis

Western blot analysis was performed on postnatal day 30 cerebellum from transgenic mice *R26:LSL-PB-ERT2^+/-^* mice crossed with *Nestin-Cre^+/-^* mice. Extracted protein was run on Novex Wedgewell 8–16% Tris-Glycine gradient gels (Thermo Fisher) then transferred to PVDF membranes. Membranes were then blocked in TTBS with 5% Skim milk (Bioshop #SKI400) for two hours and probed overnight in TTBS/1% skim milk with a 1:3000 dilution of mouse anti-α-Tubulin (Sigma Aldrich #T6199), or a 1:500 dilution of mouse anti-Erα antibody (SCBT sc-56833). Membranes were then washed in TTBS/1% skim milk, incubated with secondary antibodies (1:5000 anti-mouse IgG, HRP-linked antibody (Cell Signaling #7076S)). Finally, blots were washed in TTBS/1% skim milk, incubated in Pierce ECL Western substrate (Thermo Fisher #32209) and signal visualized on a Bio-Rad ChemiDoc system.

### Necropsy, tumor collection, and histological analysis

Experimental mice were monitored for tumor formation over a period of 365 days. When mice reached humane endpoint, they were sacrificed according to Canadian Council on Animal Care (CACC) guidelines. Upon sacrifice, cerebellar tumors were collected and divided into smaller pieces and frozen on dry ice. Samples were placed at −80 °C for long-term storage or in RNAlater (Sigma). Formalin-fixed tissue samples were paraffin-embedded by the Pathology Core at the Centre for Modeling Human Disease (CMHD) in TCP. 5 μm sections were stained with Hematoxylin and Eosin and used for histological analysis.

### Histological analysis

Formalin-fixed tissue samples were paraffin-embedded by the Pathology Core at the Centre for Modeling Human Disease (CMHD) in TCP. 5 μm sections were stained with Hematoxylin and Eosin, and used for histological analysis.

### Library preparation and sequencing

Restriction-splink PCR was performed based on previous protocols^21, 22^. In brief, mouse tumour gDNA was extracted and mechanically sheared to 300 bp fragments using a Covaris S220 sonicator. End repair was then performed using a EpiCentre End-It Kit, followed by adaptor ligation. With splinkerette-adaptors ligated at both ends, BamHI was used to digest and remove gDNA fragments. Primary and secondary PCR amplifications were performed using primers listed below for Left & Right amplification and for barcoding, respectively. Finally, purified PCR products were sent for 454 parallel sequencing at the Ontario Institute for Cancer Research. Genomic DNA libraries from LP tumours were prepared as sequenced as previously described^65^. Three libraries were prepared to identify different types of LP insertion events. To identify all LP insertions, LP insertions that had undergone PB excision, and LP insertions that had not undergone PB excision, the IR, PB, and JX libraries were prepared using primers corresponding to individual SB ITRs, both SB ITRs, and SB plus PB ITRs, respectively.

<u>Primers to generate adaptors:</u>

Linker+: 5’-GTAATACGACTCACTATAGGGCTCCGCTTAAGGGAC-3’

Linker-: 5’-Phos-GTCCCTTAAGCGGAG-C3spacer-3’

<u>Primary PCR primers for Left and Right amplification:</u> IRL (left): 5’-AAATTTGTGGAGTAGTTGAAAAACGA-3’ IRR (right): 5’-GGATTAAATGTCAGGAATTGTGAAAA-3’

Linker-A1 primer: 5’-GTAATACGACTCACTATAGGGC-3’

<u>Secondary PCR primers for barcoding:</u> IR-barcoded primer:

5’-AATGATACGGCGACCACCGAGATCTACACTCTTTCCCTACACGACGCTCTTCCGATCT(barcode)TGTATGT AAACTTCCGACTTCAACTG-3’

LinkerA2_PE (for paired-end sequencing): 5’-

CAAGCAGAAGACGGCATACGAGATCGGTCTCGGCATTCCTGCTGAACCGCTCTTCCGATCTTAGGGCTCCGCTTAAGGGAC-3’

<u>Primary PCR primers:</u>

PBL (left): 5’-CGATAAAACACATGCGTC-3’ PBR (right): 5’-CTCCAAGCGGCGACTGAG-3’

PBL2 (left): 5’-AAACCTCGATATACAGACCGAT-3’ PBR2 (right): 5’-TTACCGCATTGACAAGCACGCC-3’

Linker-A1: 5’-GTAATACGACTCACTATAGGGC-3’

<u>Secondary PCR primers (17 bp shared in both directions):</u> PB-barcoded primer: 5’-

AATGATACGGCGACCACCGAGATCTACACTCTTTCCCTACACGACGCTCTTCCGATCT(barcode)tatctttctagggttaa-3’

PB-barcoded P1: 5’-AATGATACGGCGACCACCGAGATCTACACTCTTTCCCTACACGACGCTCTTCCGATCTAAGAAGAAtatctttctagggttaa-3’

PB-barcoded P2: 5’-AATGATACGGCGACCACCGAGATCTACACTCTTTCCCTACACGACGCTCTTCCGATCTGGCAAGAAtatctttctagggttaa-3’

PB-barcoded P3: 5’-AATGATACGGCGACCACCGAGATCTACACTCTTTCCCTACACGACGCTCTTCCGATCTCCTAAGAAtatctttctagggttaa-3’

PB-barcoded P4: 5’-AATGATACGGCGACCACCGAGATCTACACTCTTTCCCTACACGACGCTCTTCCGATCTCGAGAGAAtatctttctagggttaa-3’

LinkerA2_PE (for paired end sequencing): 5’-CAAGCAGAAGACGGCATACGAGATCGGTCTCGGCATTCCTGCTGAACCGCTCTTCCGATCTTAGGGCTCCGCTTAAGGGAC-3’

<u>Primary and secondary Sleeping Beauty PCR primers:</u>

<u>Junction PCR:</u>

PBLJ 5’-gtagcattgcagtactaagc-3’

PBLJ2 5’-gtatttggtagcattgcagt-3’

PBRJ 5’-cactaagttgagtactaagc-3’

PBRJ2 5’-ggtcagaagtttacatacac-3’ PBLJX (Overlap) 5’-gcttttaaattgttaagcacaagc-3’ PBLJX2 (Overlap) 5’-ctaagcttttaaattgttaagcacaagc-3’ PBRJX (Overlap) 5’-actaagcttgtgcttaacaatt-3’ PBRJX2 (Overlap) 5’-gttgagtactaagcttgtgcttaacaat-3’ PBLJ3 5’-gcattgcagtactaagcttttaaat-3’

PBLJ4 5’-tggtagcattgcagtactaagctt-3’

PBLJ5 5’-ctcaattagtatttggtagcattgcag-3’

PBRJ3 5’-cactaagttgagtactaagcttgtg-3’

PBRJ4 5’-catacactaagttgagtactaagcttg-3’

PBRJ5 5’-gtcagaagtttacatacactaagttgag-3’

### LP read processing, alignment and analysis

Processing and gCIS analysis of Lazy Piggy transposon insertion sites were performed using a custom R script. To correct for technical alignment jitter in insertion mapping, insertions were grouped by sample and orientation (left or right), then counts were aggregated for insertions within 5bp of each other. We removed insertions mapping to non-standard or donor chromosomes, those with single read support, or those detected in *Ptch1^+/-^* control mice. A dynamic filter was used to categorize insertions as clonal or subclonal, as previously described ^23, 65^. For each library, three thresholds were calculated using the insertion data: (i) >95th percentile of reads under the negative binomial distribution fit to the number of sites with 1–3 reads, (ii) 1% of the read count of the most abundant insertion site, (iii) 0.1% of the total read number. The most stringent value was the threshold for clonal insertions and the second-most was the threshold for the clonal/subclonal category. Gene-centric common insertion site (gCIS) analysis was performed using clonal/subclonal insertions. gCIS genes were RefSeq genes (+15kb buffer) with insertions in at least 3 separate tumors and Bonferroni-corrected p value <0.05 from a Chi-square test of observed and expected insertion counts given the number of TA dinucleotide sites within the gene relative to the whole genome and the total number of insertions within each tumor. Known false positive genes *En2*, *Sfi1,* and *Foxf2* were removed since they contain sequence homology with the LP transposon. To identify tumor maintenance genes, we compared gCIS genes from PB libraries in mice with and without tamoxifen treatment. To assess robustness of tamoxifen-induced PB remobilization, Jaccard similarity scores were calculated for all pair-wise library comparisons in IR and JX libraries.

### Bulk RNAseq library preparation, data preprocessing, and analysis (Lazy Piggy tumours)

Bulk RNAseq libraries were prepared and sequenced as previously described^66^ using the Miseq system with 5 samples pooled per lane and mean 50 million reads per sample. After raw FASTQ quality check with FastQC, reads were aligned using STAR (2.5.4b) to mouse genome mm9 using the annotation file Mus musculus NCBIM37v67 (downloaded from Ensembl) with masking for the *En2* gene, whose splice acceptor sequence is contained in the LP transposon^67^. The “ReadsPerGene” raw counts from STAR were used for differential expression analysis with DESeq2 (1.34.0) using genes with non-zero counts in at least two samples per tamoxifen treatment group^68^. Dimensionality reduction by UMAP^69^ was then performed to visualize the distribution of tamoxifen receipt status across clusters following variance-stabilizing transformation of raw counts in DESeq2.

### Bulk RNAseq analysis (Human MB and normal cerebellum)

Normalized counts were generated from published human MB and normal cerebellar bulk RNAseq data as described previously^58, 70, 71^. DESeq2 (1.34.0) was used for differential expression analysis^68^.

### Single cell RNAseq analysis

Normalized counts for human SHH MB samples were downloaded from GEO^34^ and UCSC Cell Browser^35^ and prepared as Seurat (4.1.0) objects. UMAP embeddings for the data from Riemondy et al. were directly downloaded. UMAP embeddings for Hovestadt et al. were calculated following the Seurat SCTransform() vignette and batch effects corrected using {harmony} (0.1.0) with all default settings. Joint densities were plotted to visualize co-expression of selected genes using {Nebulosa} (1.4.0).

### Mouse studies

All procedures were performed in compliance with the Animals for Research Act of Ontario and the Guidelines of the Canadian Council on Animal Care. The Centre for Phenogenomics (TCP) Animal Care Committee reviewed and approved our protocol 19-0288H. In all cases, mice were maintained on a 12-hour light/dark cycle with free access to food and water. For all studies, mice of either sex were used, and randomly allocated to experimental groups. *Ptch1^+/-^*^15^, *Math1-Cre*^72^, *Rosa26-LSL-SmoM2-YFP* (*SmoM2*)^73^, *Kcnb2^-/-^* (*Kcnb2^tm1Lex^*)^74^, and *Nestin-Cre*^75^ mice were previously described. All mice were bred and genotyped as recommended by Jackson Laboratories. *Math1-Cre; SmoM2* mice develop SHH MB due to expression of SmoM2—a constitutively active form of the SHH pathway receptor Smoothened—in cerebellar granule neuron precursors (CGNPs), achieved by CGNP-specific driver *Math1-Cre*^17, 18^. *Ptch1^+/-^* mice develop SHH MB due to loss of one allele of the SHH pathway inhibitor *Ptch1*^15^ and subsequent loss-of-heterozygosity, which drives constitutive SHH signaling in CGNPs.

For anti-Hedgehog therapy, 50 mg/kg vismodegib (GDC-0449, Selleck Chemical) was administered three times (P18, 19, 20) in 4.76% DMSO 0.5% methylcellulose 0.2% Tween 80 buffer by intraperitoneal injection. For single-dose BrdU labeling, mice were injected intraperitoneally with 50 mg/kg BrdU (Roche) in PBS. Tamoxifen-citrate mixture at 400 mg/kg was incorporated to the standard rodent diet premixed with ∼5% sucrose as a palatability enhancer (Harlan Laboratories Teklad Diets). Tamoxifen chow was introduced once tumour-induced cranial bulge visualized, typically 45–60 days postnatal.

### Mouse Sox2^+^ MB cell isolation and culture

Mouse Sox2^+^ MB cells were isolated as previously described^37^. Briefly, brain tumors from P7 *Math1-Cre; SmoM2* mice were dissociated by repetitive pipetting using ice-cold PBS without Mg^2+^ and Ca^2+^, followed by treatment using 50% Accutase (Stemcell Technologies) diluted in PBS. Dissociated cells were cultured on plates coated with poly-L-ornithine (Sigma-Aldrich) and laminin (Sigma-Aldrich), using Neurocult NS-A Basal media (StemCell Technologies) supplemented with 2 mM L-glutamine, N2, B27, 75 μg/ml BSA, 2 μg/ml Heparin, 10 ng/ml basic FGF and 10 ng/ml human EGF without addition of serum. All cell lines were regularly checked for mycoplasma infections.

For cell counting assays, cells were plated at a density of 2000 cells per well in poly-L-ornithine- and laminin-coated 12 well plates. At 2, 4, or 6 days after seeding, cells were resuspended using Accutase and incubated in isotonic solution. Cell count was acquired using a Multisizer 4 Coulter Counter (Beckman-Coulter) using standard protocols and a threshold of 12 to 30 μm.

### Immunocytochemistry

Immunocytochemistry was performed on cultured cells as previously described^76^. Briefly, cells on glass coverslips were fixed for 15 minutes with 4% PFA and then permeabilized with 0.1% Triton X-100 in PBS (PBST). Cells were subsequently blocked with 10% normal goat serum in PBST for one hour at room temperature and incubated with primary antibodies in blocking solution overnight at 4° C, followed by incubation with fluorophore-conjugated secondary antibodies (1:400-1000) and 1 μg/ml DAPI (Sigma-Aldrich) for one hour at room temperature. For F-actin staining, cells were stained with 1:500 Alexa Fluor 488 Phalloidin (Invitrogen). Coverslips were mounted onto glass slides using Prolong Gold (Invitrogen). The primary antibodies include: chicken anti-GFP (Antibodies Incorporated #GFP-1020, 1:1000), mouse anti-α-Tubulin (Sigma-Aldrich #T6199, 1:1000), mouse anti-BrdU (DSHB #G3G4, 1:1000), mouse anti-Caveolin-1 (Novus #NB100-615, 1:200), rabbit anti-Caveolin-1 (Cell Signaling Technology #3267, 1:400), rabbit anti-Clathrin Heavy Chain (Cell Signaling Technology #4796, 1:100), rabbit anti-pEGFR (Cell Signaling Technology #3777, 1:800), rabbit anti-pEGFR (abcam #ab40815, 1:250), rabbit anti-pERM (Cell Signaling Technology #3726, 1:200), rabbit anti-Rab5 (Cell Signaling Technology #3547, 1:400). Images were acquired using a Leica SP8 Lightning Confocal DMI6000 microscope. Images were analyzed using Imaris software.

### Tissue preparation and immunohistochemistry

Postnatal (P7 and P21) mice were transcardially perfused with ice-cold PBS, followed by 4% PFA. Brains were removed and fixed in 4% PFA overnight upon collection. Brains were then cryopreserved in 30% sucrose for 48-72 hours, mounted in O.C.T. compound (Tissue-tek), and cryo-sectioned at 10-12 μm. For immunohistochemistry, frozen sections were dried at room temperature for 30 minutes and rehydrated in PBS with 0.1% Tween (PBSTw). Antigen retrieval was performed in 10mM citrate buffer, pH 6.0, for 20 minutes at 95° C. Sections were blocked with 10% normal goat serum in PBSTw for one hour at room temperature and incubated with primary antibodies in blocking solution overnight at 4° C, followed by incubation with fluorophore-conjugated secondary antibodies (1:200-500) and 1 μg/ml DAPI (Sigma-Aldrich) for one hour at room temperature. Sections were then mounted with glass coverslips using Aquamount (Fisher Scientific). The primary antibodies include: mouse anti-BrdU (DSHB #G3G4, 1:100), rat anti-BrdU (abcam #6326, 1:500), mouse anti-KCNB2 (NeuroMab #75-369, 1:100), mouse anti-PCNA (Santa Cruz Biotechnology #sc-56, 1:200), mouse anti-SOX2 (abcam #ab79351, 1:100), rabbit anti-SOX2 (abcam #ab97959, 1:200), rabbit anti-Caveolin-1 (Cell Signaling Technology #3267, 1:200), rabbit anti-DCX (abcam #ab18723, 1:200), rabbit anti-Ki67 (abcam #ab15580 1:200), rat anti-Ki67 (Ebioscience #14-5698-82, 1:200), rabbit anti-NeuN (abcam #ab104225, 1:200), rabbit anti-pEGFR (abcam #ab40815, 1:250), rabbit anti-pERK1/2 (Cell Signaling Technology #4370, 1:200), rabbit anti-pERM (Cell Signaling Technology #3726, 1:200). Cell death was determined using TUNEL (Cat. # S7110, Sigma-Aldrich) according to the manufacturer’s instructions. Images were acquired using a Leica SP8 confocal microscope or a Quorum spinning disc confocal microscope. Images were analyzed using Imaris software. Hematoxylin and eosin staining was performed on paraffin-embedded sections and imaged using a 3D Histech Pannoramic 250 Flash II Slide Scanner.

### *In vitro* limiting dilution assay

Cells were plated in serial dilutions on non-adherent 96-well plates and in six biological replicates under stem cell conditions. Serial dilutions ranged from 2000 cells to 3 cells per well. After 7 days of plating, each well was scored for negative spheres. Data was plotted and tested for inequality in frequency between multiple groups and tested for adequacy of the single-hit model using Extreme Limiting Dilution Analysis (ELDA) software.

### Atomic force microscopy

Force-displacement data were collected at room temperature using an AFM (Bioscope Catalyst, Santa Barbara, CA) mounted on an inverted microscope (Nikon Eclipse Ti2). Force-displacement-speed data were measured at the cell center. Measurement of cells in each petri dish was completed within 20 mins after being taken out of the incubator. The AFM probe with a spherical tip was used to measure the cell stiffness, using the Hertz model to calculate the cell stiffness value from the force-displacement data. The AFM probe with a needle shape tip was used to indent and penetrate the cell membrane, for measuring the cell membrane tension. The cell membrane tension results were calculated based on the force-displacement data before the membrane rupture was observed (larger than 100 pN) and using the mechanics model based on previous studies ^77^. The AFM probes used for cell stiffness measurement experiments were biosphere B100-CONT (Nanoandmore, USA), with a nominal spring constant of 0.2 N/m. The AFM probes used for cell membrane tension experiments were Focused-Ion-Beam modified probe MSNL-10 (Brucker, USA), with a nominal spring constant of 0.03 N/m ^78^. The spring constant of each probe was calibrated using thermal spectroscopy (Nanoscope 8.10). The loading speeds were set to be 10 μm/s to minimize the effects from the viscoelastic properties of the cell. Data analysis for quantifying cell stiffness and cell membrane tension was conducted in MATLAB. The code of data analysis for rejecting the non-rupture case is available for download at https://github.com/XianShawn/Nuclear_Mechanics.

### Micropipette aspiration

The micropipette was fabricated from a glass capillary using a commercialized micropipette puller system (Model P-97, Sutter Instrument). The inner diameter of the micropipette tip is 2 μm. To make the tip horizontal under the microscope for clear aspiration observation, the tip was bent by 45° using a Microforge (Model MF-l, TPI Instruments). The pressure applied to the tip of the micropipette was controlled by a pneumatic microinjection pump (Digital Microinjector from Sutter Instrument). Before aspiration, a positive pressure was applied to the micropipette tip to balance the capillary force. During aspiration, the micropipette tip was gently brought into contact with the cell surface and the pressure applied to the tip was reduced by a set amount. The entire aspiration process was recorded by a camera (scA1300-32gm, Basler) at 33 frames per second. The aspiration length and speed of the cell inside the micropipette was measured manually. The inner cell pressure was calculated using the standard linear solid model.

### Electrophysiology

Cells were cultured on laminin-coated plastic coverslips for 48-72 hours. Coverslips were transferred to a recording chamber filled with bath solution. The bath solution consisted of (in mM) 118 NaCl, 3 KCl, 2.5 CaCl2, 1.5 MgCl2, 10 glucose, and 10 HEPES (PH adjusted to 7.4 with NaOH). Patch pipettes (borosilicate glass) for recording, with resistance of around 4 MΩ, were filled with intracellular solution consisting of 125 mM KCl, 11 mM EGTA, 1 mM CaCl2, 1.5 mM MgCl2, and 10 mM HEPES (PH adjusted to 7.2 with KOH). Whole-cell currents were recorded using an Axopatch 700B amplifier (Molecular Devices). All experiments were performed at room temperature. Pipette and whole cell capacitance were compensated. The voltage protocol consisted of 200 ms pulses from -80 mV to +80 mV (20 mV voltage steps). Data were acquired online, filtered at 4 kHz, digitized at 10 kHz, and analyzed offline using pClamp10 (Molecular Devices). Leak currents before voltage stimulations were subtracted off-line. I-V curves were generated by plotting peak current amplitude at different voltages. Data were quantified and graphed using GraphPad Prism.

### Statistical analyses

No statistical methods were used to pre-determine sample sizes. The statistical analyses were performed after data collection without interim data analysis. No data points were excluded. Two-tailed Student’s t-test was performed for comparison between two groups of samples. Two-Way ANOVA analyses were used to assess significance of multiple data points. The Kaplan–Meier estimator was used to generate survival curves using the R package {survival}^79^. Differences between survival curves were calculated using a log-rank test. All data were collected and processed randomly. All data are expressed as mean ± SEM. We considered a *P* value less than 0.05 to be statistically significant.

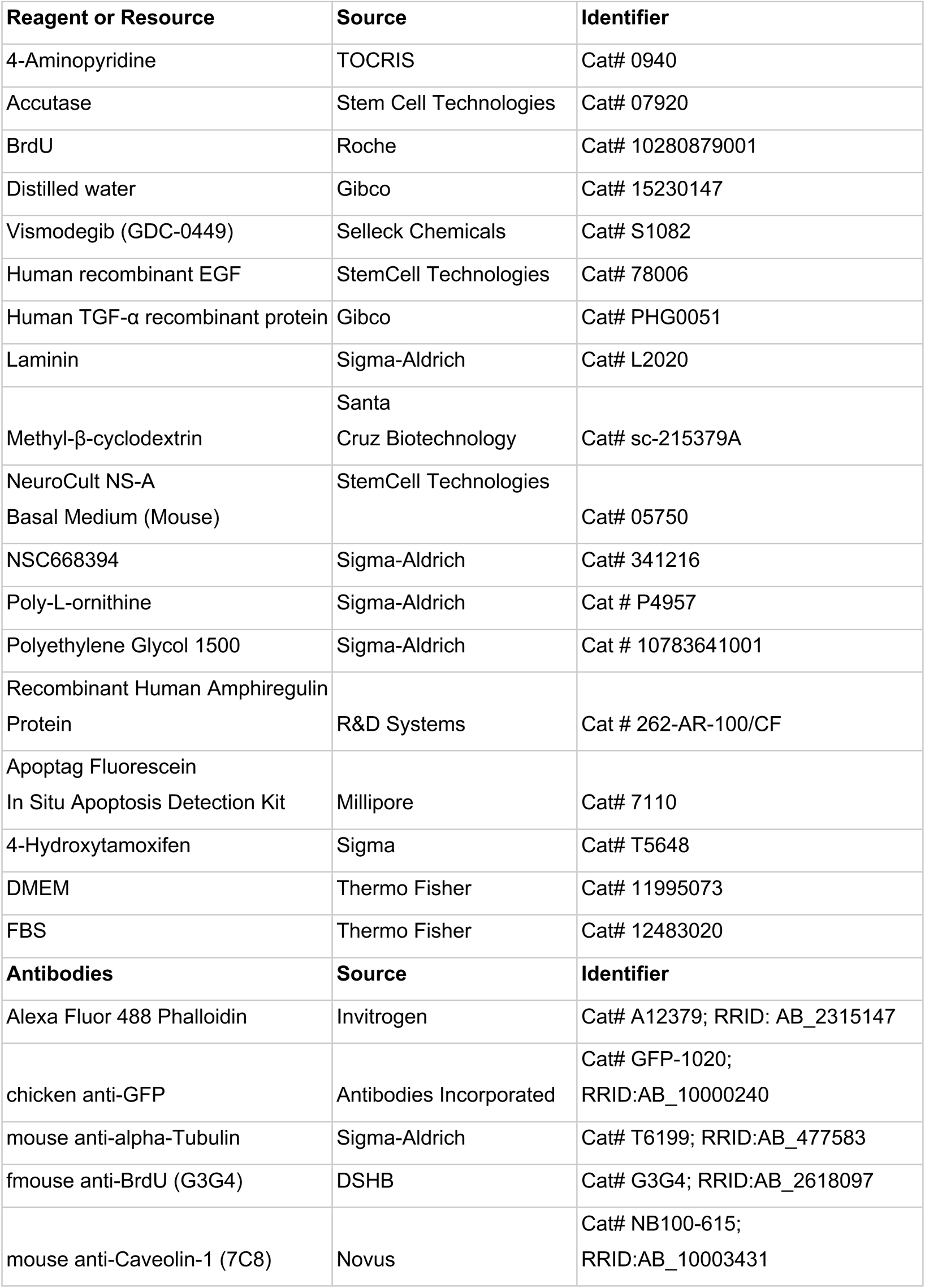
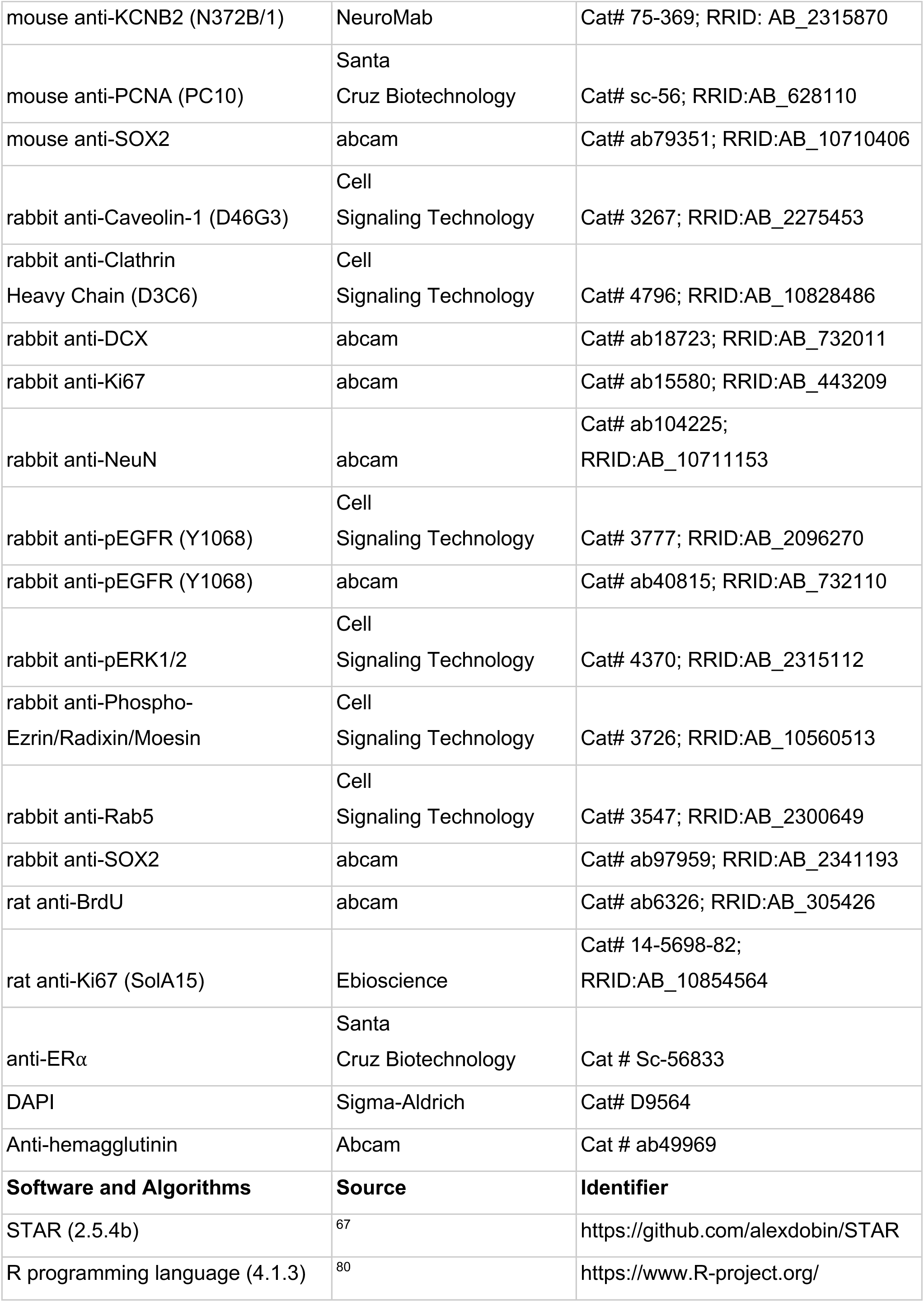
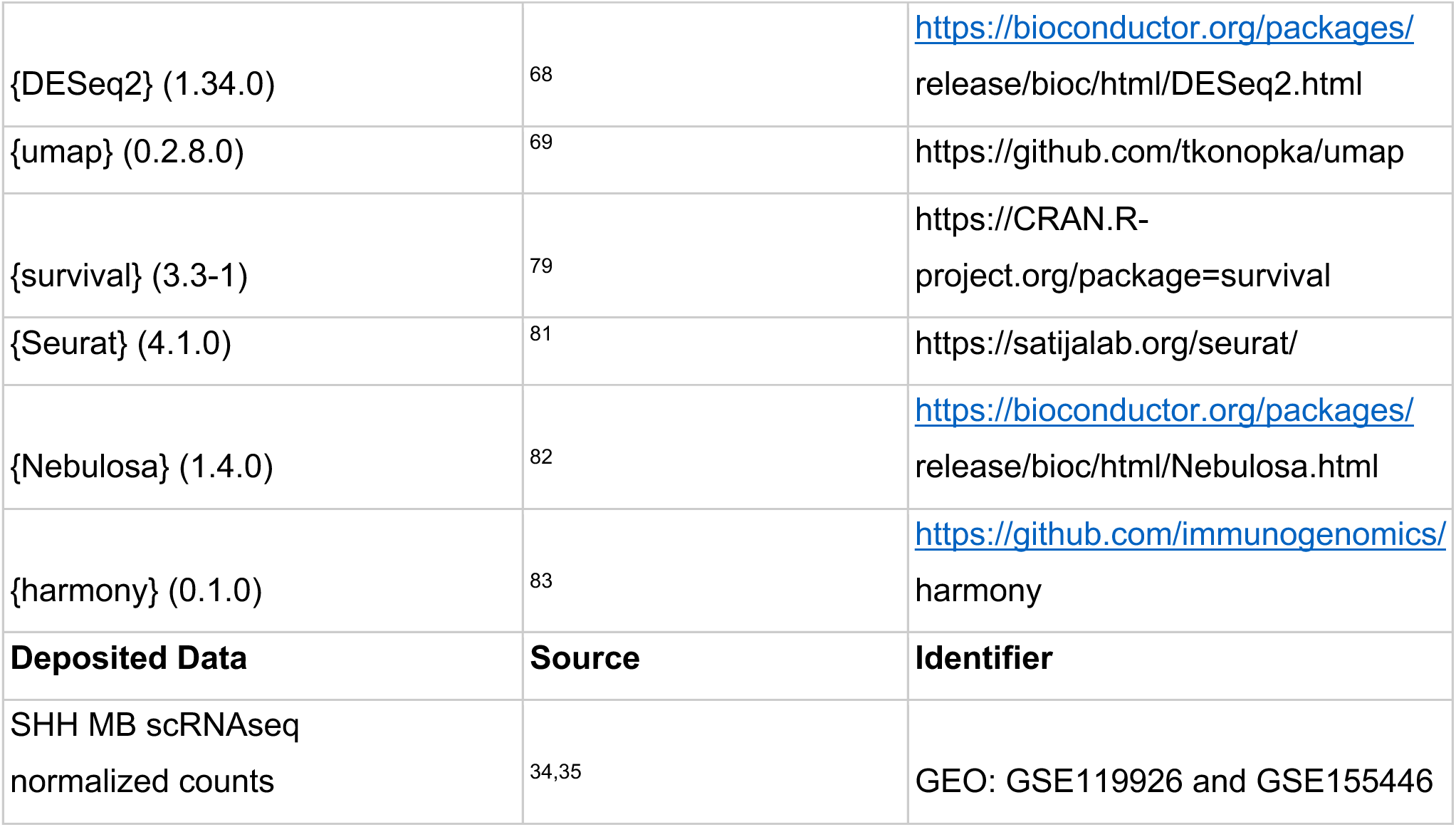
Key Resources Table

## RESOURCE AVAILABILITY

### Lead Contact

For all additional information and request for resources and reagents should be directed to and will be fulfilled by the Lead Contacts, Dr. Xi Huang and Dr. Michael Taylor.

### Materials Availability

All unique materials and reagents generated in this study can be obtained from the Lead Contact.

### Code Availability

Custom code used in the analyses is deposited on GitHub (https://github.com/anderswe/lazy_piggy).

## Notes

### Competing Interest Statement

The authors have declared no competing interest.

### Summary of Updates

Additional data added to Figure 6 and Supplemental Figure 7.

